# Enhancing cancer targeting of γ9δ2TCR through modified NKG2D co-stimulation

**DOI:** 10.1101/2021.01.06.424553

**Authors:** Patricia Hernández-López, Eline van Diest, Inez Johanna, Sabine Heijhuurs, Trudy Straetemans, Zsolt Sebestyén, Dennis X. Beringer, Jürgen Kuball

## Abstract

Despite the ability of γδT cells to mediate tumor killing independently of MHC recognition, all the clinical trials that have been carried out using these cells showed low response rate in patients, in part due to its poor proliferation ability. Recently, a new generation of CAR-T cells called αβT cells engineered to express a defined γδTCR (TEG) has been developed. TEGs are αβT cells engineered to express a defined γδTCR. These cells are able to mediate effective antitumor reactivity without showing any reactivity towards healthy tissue, and combine the best qualities of both αβT and γδT cells. In fact, the high affinity γ9δ2TCR clone 5 has recently been selected within the TEG format as a clinical candidate (TEG001). Here we present a strategy to improve the antitumor activity of TEG001 by co-expressing an activating chimeric co-receptor together with γδTCR-Cl5.Therefore, we developed three different co-receptors by fusing the extracellular domain of the activating cell surface receptor NKG2D, that is able to bind stress induced ligands typically expressed on tumor cells, to the cytoplasmic signaling domains of the T cell costimulatory proteins ICOS, CD28 and 4-1BB. We determined that introduction of the chimeric co-receptors NKG2D-CD28_wt_ and NKG2D-4-1BB_CD28TM_ improved the activity of TEG001 against tumors that were recognized by γδTCR-Cl5 and expressed NKG2D ligands, but did not affect tumors that either were not recognized by γδTCR-Cl5 or did not express NKG2D ligands. This ‘chimeric co-receptors’ approach open a wide range of opportunities that lead to a next generation of TEGs.

## Introduction

It has been demonstrated that TEGs (αβT cells engineered to express a defined γδTCR) mediate effective antitumor reactivity without showing any reactivity towards healthy tissue and present a novel class of CAR T which has the potential to target many different tumor types [1]. For CAR T, despite its huge success in the clinic further improvements have been suggested such as dual receptor signaling in order to increase safety and or efficacy [2]. In this light we explored whether either changing signaling domains of the γδTCR or providing additional costimulatory signals could improve performance of TEGs. Different signaling capacities of αβTCR and γδTCR have been reported to impact long term memory formation of γδT cells [3] CD8^+^ TEG activity has been reported to be partially supported through endogenous NKG2D [4]. NKG2D is an activating cell surface receptor that is able to bind to 8 different several stress induced ligands (MICA/B and ULBP1-6) that are overexpressed in many tumor cells but absent or present at low levels in healthy cells [5]. NKG2D-NKG2D ligand interaction has been shown alone to be sufficient to induce killing of tumor cells through γδT cells [6] and NK cells [5]. As consequence NKG2D CAR αβT cells have been explored in clinical trials but showed no substantial efficacy in patients suffering from acute myeloid leukemia [7]. Reasons of failure might be that NKG2D alone without additional coreceptor support does not exploit its full activity or that the used design which partially depended on DAP10 signaling rapidly exhausts and contributed to education of engineered immune cells. γδT cells and NK cells have been shown to quickly adopt to new environments and get educated over time, thus tolerant [8, 9]. Within this context we explored whether adding NKG2D to CD4+ and CD8+ T cells in its wild-type version or with altered signaling domains can further enhance the promising and broad anti-tumor activity of a γ9δ2TCR. As additional signaling domains different costimulatory proteins (ICOS, CD28 and 4-1BB) that are typically expressed on T cells and also used for CAR T engineering [2] have been explored.

## Material and Methods

### Antibodies

The following antibodies were used: anti-CD8a-PerCP-Cy5.5 (1:100; clone RPA-T8; 301032), CD4-AF700 (1:40; clone RPA-T4; 300526), αβTCR-PE-Cy5 (1:80; clone IP26; 11-9986-42) from Biolegend; γδTCR-PE-Cy7 (1:20; clone IMMU510; B10247) from Beckman Coulter;NKG2D-PE (1:20; clone 1D11; 320806), NKG2D-BV650 (1:40; clone 1D11; 563408) and γδTCR-APC (1:5; clone B1; 555718) from BD Biosciences.

### Cell lines and cell culture

Daudi, K562, HL60, Jurkat, RPMI-8226 and Phoenix-Ampho cells were obtained from ATCC. Phoenix-Ampho cells were cultured in DMEM supplemented with 1% Pen/Strep (Invitrogen) and 10% FCS (Bodinco, Alkmaar, The Netherlands). All other cell lines were cultured in RPMI with 1% Pen/Strep and 10% FCS. Primary fresh PBMCs were isolated by Ficoll-Paque (GE Healthcare, Eindhoven, The Netherlands) from buffy coats supplied by Sanquin Blood Bank (Amsterdam, The Netherlands).

### Construction of chimeric NKG2D receptors

Six chimeric receptors were constructed (for sequences see Appendix I). cDNAs for co-receptors were synthetized by BaseClear (Leiden, Netherlands). Type II co-receptors were created using overlap extension PCR. For the first reaction DNA coding for cytoplasmic signaling domains of ICOS and CD28, both extended with 18 nucleotides of the NKG2D transmembrane sequence were amplified using the following primers; for ICOS “ICOScyto_FW1“ and “NKG2D_OlEx_RV1”, and for CD28 “CD28cyto_FW1 “ and “NKG2D_OlEx_RV1”. For the second reaction the DNA for “4-1BBcyto-NKG2D tm + ec” was used as template for the transmembrane and extracellular domain of NKG2D and was amplified using the following primers “NKG2D_FW2” and “NKG2D_BamHI_RV2”. In reaction 3 the products of reaction 1 were fused to the product of reaction 2 using “ICOScyto_FW1“ and “NKG2D_BamHI_RV2” for ICOScyto-NKG2D, and “CD28cyto_FW1 “ and “NKG2D_BamHI_RV2” for CD28cyto-NKG2D. All type I and II constructs were subcloned into pBullet using NcoI and BamHI as restriction sites.

In similar fashion as the type II constructs the transmembrane and linker domains of the type I co-receptors NKG2D-ICOS_wt_ and NKG2D-4-1BB_wt_ were replaced by the transmembrane and linker domains of NKG2D-CD28_wt_ chimera using overlap extension PCR. In reaction 1 NKG2D-CD28linker-tm was amplified using primers “P2A_FW2 “ and “CD28tm_RV2” and in reaction 2 the signaling domains of ICOS and 4-1BB were amplified using primers “ICOS_FW1” or “41BB_FW1” in combination with “pmp71_RV”. In reaction 3 the PCR products of reaction 1 and 2 were fused using primers “P2A_FW2“ and “pmp71_RV”. They were subcloned into pMP71 already containing γδTCR-Cl5, using XhoI and HindIII. All restriction enzymes were supplied by NEB (Massachusetts, USA).

### Retroviral transduction of αβ T cells and cell lines

Briefly, packaging cells (Phoenix-Ampho) were transfected with helper constructs gag-pol (pHIT60), env (pCOLT-GALV) and pMP71 or pBullet retroviral vectors containing genes codifying for the different proteins. In the case of human PBMCs, they were pre-activated with anti-CD3 (30 ng/mL; Orthoclone OKT3; Janssen-Cilag) and IL-2 (50 IU/mL; Proleukin, Novartis). Both, PBMCs and cell lines, were transduced twice with viral supernatant within 48 or 3 hours respectively, in presence of 6 mg/mL polybrene (Sigma-Aldrich). For PMBCs 50 IU/mL of IL-2 was added. TCR-transduced T cells were expanded by stimulation with anti-CD3/CD28 Dynabeads (500,000 beads/10^6^ cells; Life Technologies) and IL-2 (50 IU/mL). Thereafter, TCR-transduced T cells were depleted of the non-engineered T cells.

### Depletion of non-engineered T cells

αβT cells transduced with γδTCR-Cl5 either alone or together with NKG2D wild type or the different NKG2D chimeras were incubated with a biotin-labeled anti-αβTCR antibody (clone BW242/412; Miltenyi Biotec, Bergisch Gladbach, Germany) and subsequently incubated with an anti-biotin antibody coupled to magnetic beads (anti-biotin MicroBeads; Miltenyi Biotec). Thereafter, the cell suspension was loaded onto an LD column and αβTCR+ T cells were depleted by MACS cell separation per the manufacturer’s protocol (Miltenyi Biotec). After depletion, TEGs were expanded using T cell REP.

### Selection of engineered T cells

After αβ-depletion, T cells were selected using human CD4 microbeads and MS columns (Miltenyi Biotec). Procedure was carried out according to manufacturer’s protocol. For Jurkats, CD3 selection was performed after being transduced with γδTCR-Cl5 using human CD3 microbeads (Miltenyi Biotec). Moreover, another selection was carried out in these cells after the transduction of the different NKG2D-chimeras. First, cells were incubated at room temperature for 30 min using a 1:20 dilution of anti-NKG2D-PE in MACs buffer (PBS, 2% FCS, 2mM EDTA). Cells were washed once with MACs buffer. After washing, cells expressing NKG2D were selected using anti-PE microbeads (Miltenyi Biotec) following manufacturer’s protocol.

### NKG2D ligand staining

The expression of NKG2D ligands in tumor cell lines was assessed using Recombinant Human NKG2D Fc Chimera Protein (R&D systems, Abingdon, UK). 10^5^ tumor cells were incubated either with 0.5 μg of NKG2D Fc recombinant protein or IgG1-Fc during 30 min. Cells were washed with FACs buffer (1% BSA, 1% Na^+^azide) and secondary antibody IgG-PE (Southern Biotech, Alabama, USA) was added in a 1:200 dilution. Cells were fixed using 1% PFA in PBS. Samples were measured on a BD LSRFortessa and FACSDiva (BD) software was used for data analysis.

### Functional T cell assays

To assess T cell activation by surface recaptor expression like CD69 a FACS-based assay was used. To allow better differentiation on FACs, target cells were labeled using Cell Trace Violet Proliferation Kit (Thermo Fisher Scientific, Massachusetts, USA). T cells were resuspended at 1 × 10^6^ cells/ml in 2 μM Cell Trace Violet in PBS solution. Cells were washed two times with complete RPMI medium and resuspended in culture medium. After labelling, 10^5^ transduced Jurkats and 2 × 10^5^ target cells were co-cultured for 18 hours in round-bottom 96-well plates in presence or absence of 100 μM pamidronate. After incubation, cells were harvested and analyzed by flow cytometry to check CD69 expression. For cytokine detection 5 × 10^4^ effector T cells and 5 × 10^4^ target cells were co-cultured for 18 hours in round-bottom 96-well plates in presence or absence of pamidronate. After incubation, supernatants were collected and either frozen or used to detect IFNγ levels straight away. ELISA was performed using IFN gamma Human Uncoated ELISA Kit (Thermo Fisher Scientific, Massachusetts, USA). For assessing cytotoxicity 5 × 10^3^ RPMI 8226 cells expressing luciferase-GFP were co-cultured with the different effector T cells at several effector: target ratios (30:1, 10:1, 3:1 or 1:1) for 18 hours in round-bottom 96-well plates. Co-cultures were done in presence or absence of 10 μM pamidronate. RPMI-8226 luciferase-GFP transduced tumor cells were used as targets. After incubation, luciferin was added at 12,5 ug/ml to each well and signal was measured on Softmax pro machine. Specific lysis was calculated using the formula % specific lysis = 100 × [(experimental data – spontaneous cell death)/(maximum cell death − spontaneous cell death)]. When this calculation provided a negative value, 0% was assigned as the result. In order to assess proliferation T cells were resuspended at 1 × 10^6^ cells/ml using a 2 μM solution of CellTrace™ Violet Cell Proliferation Kit (Thermo Fisher Scientific, Massachusetts, USA) in PBS. The cell suspension was incubated for 20 min at 37ºC. Cells were washed two times with complete RPMI medium and resuspended in culture medium. After labelling, 2,5 × 10^5^ effector T cells were co-cultured together with 2,5 x 10^5^ tumor cells in 48-well plates for 6 days. 100 μM pamidronate was added to cultures boost recognition. On day 4, medium was replaced. On day 6, cells were analyzed by flow cytometry.

## Results

### Design and expression of αβ-γδTCR chimera

As signaling of αβTCR and γδTCR differ and this could impact as reported long term memory formation [3] we explored stability and function of different designs of αβ-γδ TCR chimeras. A first version of three different αβ-γδ TCR chimeras was generated (Supplementary figure 1A), but only one of them showed stable expression in Jurkat 76 (Supplementary Figure 1B). However, it was not able to induce activation of Jurkats-76 (Supplementary figure 1C). We hypothesized that this lack of activity could be due to the existing differences in the variable-constant interphases of γδ and αβ TCRs. Therefore, we generated three new designs modifying those interphases (Supplementary figure 2A). Moreover, interphase between both constant domains was modified to increase pairing and mouse residues were added to the β-constant chain to improve detection. In this case two of them were expressed (Supplementary figure 2B), but again all chimeric TCR variants were not able to induce activation of Jurkat-76 (Supplementary figure 2C) and therefore this option was not further explored.

### Design and expression of NKG2D chimeras

In order to assess whether adding NKG2D co-receptor signaling to CD4+ and CD8+ TEGs can further improve activity in addition to the wild-type version of NKG2D further constructs were engineered by fusing the extracellular domain of NKG2D to the cytoplasmic signaling domains of three different costimulatory proteins (ICOS, CD28 and 4-1BB). The natural orientation of NKG2D differs however from the other three costimulatory proteins. ICOS, CD28 and 4-1BB are type I transmembrane proteins (N-terminal is outside of the cell), while NKG2D is a type II membrane protein (N-terminal domain is inside of the cell) that does not contain any known signaling motif within its intracellular domain. Therefore, NKG2D associates in natural γδT cells or NK cells with the adaptor protein DAP10 via charged residues in its transmembrane domain [8]. Taking all this into account, two different types of chimeras were designed and generated. Firstly, in line with the natural design of NKG2D, a type II membrane protein was designed (Figure 1A, depicted as “type II design”) by fusing the transmembrane domain of NKG2D to the cytoplasmic domains of ICOS, CD28 or 4-1BB. In addition, a type I design was created by fusing extracellular domains of NKG2D to the cytoplasmic domains of ICOS, CD28 or 4-1BB (Figure 1B). Consequently, linker and transmembrane domains were different for all constructs for the “type I design”, while in the “type II design” only the signaling domain differed.

**Figure 1.**
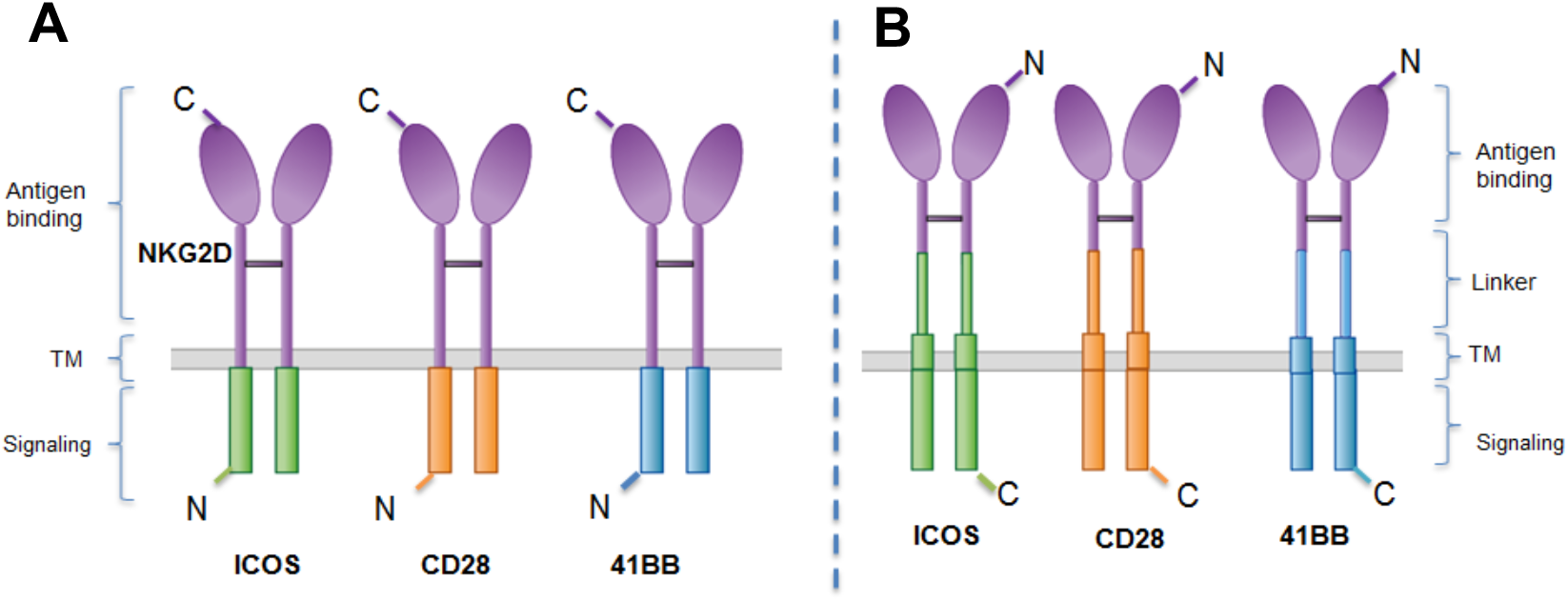
Schematic diagram of type IIand type I chimeric NKG2D co-receptors. NKG2D, ICOS, CD28 and 4-1BB regions are colored in purple, green, orange and blue, respectively. (**A**) “Type II design”. The natural NKG2D type II structure is preserved and signaling domains are added. (**B**) “Type I design” Type I transmembrane chimeric co-receptors design is build on the type I structure on the natural co-receptors ICOS, CD28 and 41BB.

To test if the design of all chimeras allows correct folding and expression, Jurkat 76 cells, that had been previously transduced and selected to express the γδTCR-Cl5, were used to transduce the different NKG2D chimera constructs. Expression of both, NKG2D-chimeras and γδTCR-Cl5, was assessed by FACS (Figure 2A). Interestingly, large differences in surface expression of the chimeric co-receptors were observed between type II and type I chimeras. For the type II designs only one of the constructs (NKG2D-4-1BB) was marginally expressed. By contrast, all type I designs were expressed in Jurkat-76. However, expression levels differed between the different type I chimeras, NKG2D-CD28_wt_ was the one that showed best expression. The observed difference in expression was also still observed when cells have been purified by MACS sort using a NKG2D antibody (Figure 2B), suggesting a different expression strength presumably mediated through differences in linker and transmembrane domains between all type I design.

**Figure 2.**
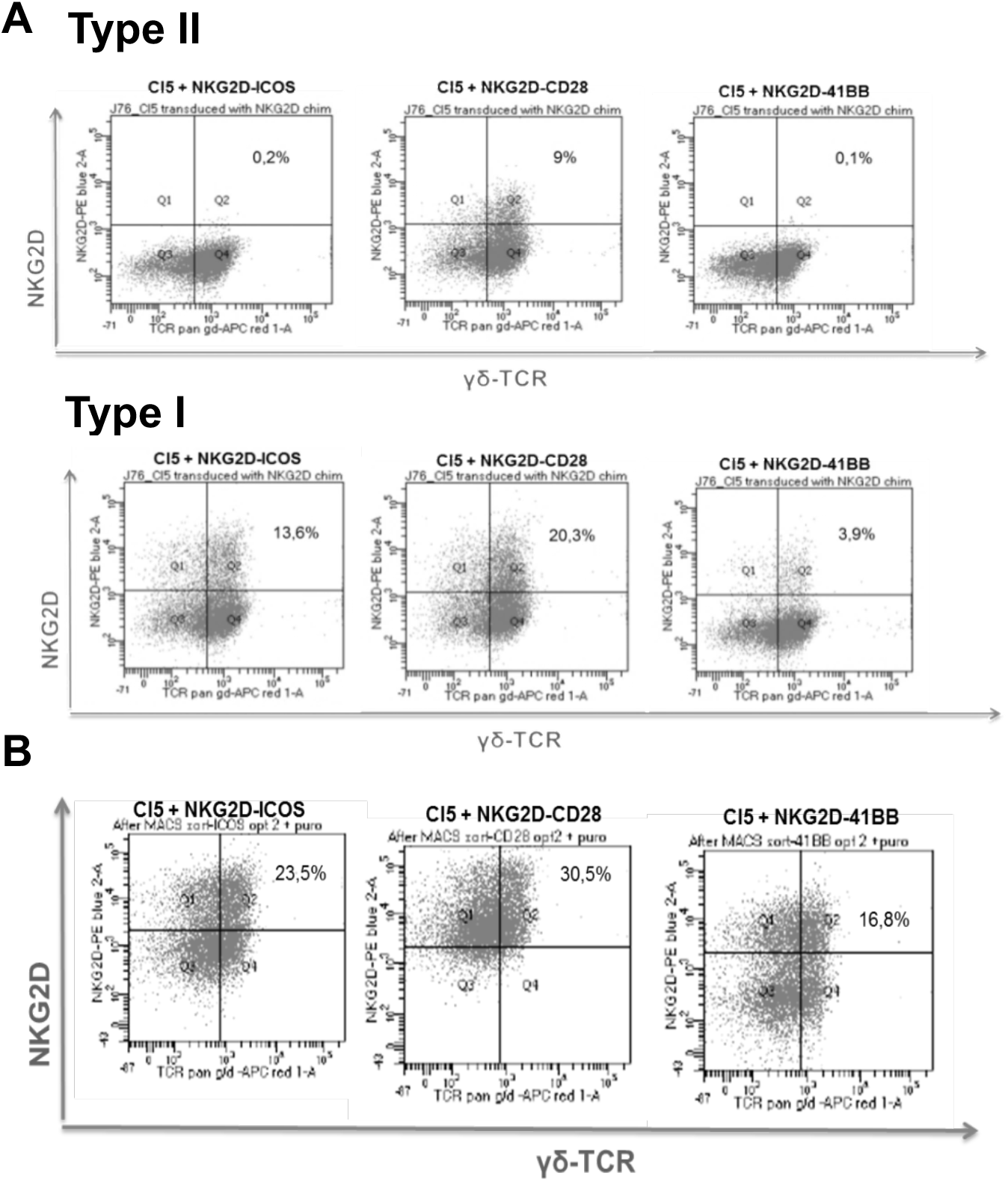
Surface expression of NKG2D chimeric co-receptors and γδTCR-Cl5 in Jurkat-76. (**A**) Surface expression of γδTCR-Cl5 and type II and type I chimeras (**B**) Surface expression of γδTCR-Cl5 and type I NKG2D chimeras after MACS sort selection using anti-NKG2D-PE and PE-microbeads.

### Introduction of NKG2D-CD28 chimera in Jurkat-76 transduced with γδTCR-Cl5 increases percentage of CD69 expressing cells

To asses if the chimeras were able to increase the activity of the Jurkats-76 transduced with γδTCR-Cl5 and the type I NKG2D chimeras, we performed a CD69 assay. Firstly, we typed target cells for NKG2D ligand expression. Target cell lines were selected according to their susceptibility to be recognized by γδTCR-Cl5 to serve as susceptible (K562, RPMI 8226, Daudi) or resistant (HL60) to TEG therapies [4, 10–13]. NKG2D ligand expression has been reported for most of these cell lines for individual ligands by us [4]. Recognition of K562, RPMI 8226, Daudi and HL60 cell lines by γδTCR-Cl5 has been previously reported and assessed in the lab. However as 8 different ligands can be bound NKG2D and we were interested in the additive effect of ligand expression to NKG2D binding, we determined the NKG2D-ligand expression by FACS using a NKG2D-Fc fusion protein. According to its susceptibility to be recognized by γδTCR-Cl5 and the expression of NKG2D ligands, we were able then to distinguish three types of cell lines, one that was recognized by γδTCR-Cl5 but expressed limited levels of NKG2D ligands (Daudi), a second one that was targeted by γδTCR-Cl5 and expressed high levels of NKG2D ligands (K562 and RPMI 8226) and a third one that was not recognized by γδTCR-Cl5 but expressed high levels of NKG2D ligands (HL60) (Figure 3).

**Figure 3.**
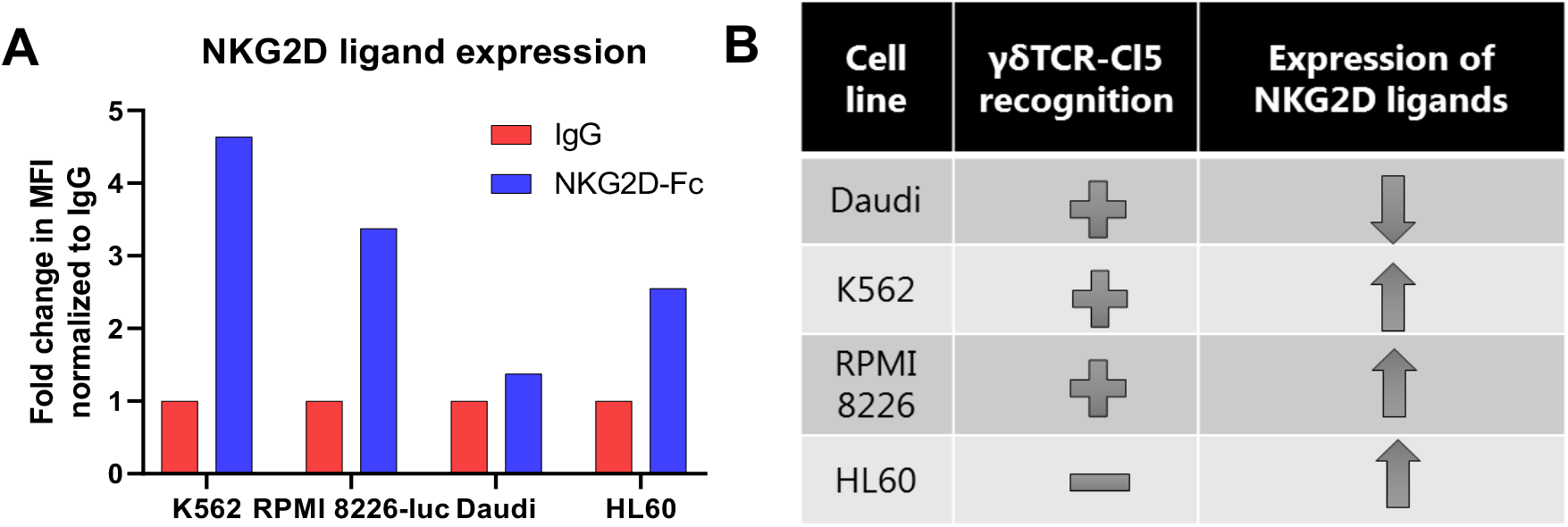
NKG2D ligand expression. (**A**) Surface expression of NKG2D ligands in Daudi, K562, RPMI 8226 and HL60, as assesed by flow cytometry (**B**) Summary of gdTCR-Cl5 recognition and NKG2D ligand expression in Daudi, K562, RPMI 8226 and HL60.

To assess activity of engineered Jurkat cells with or without engineered type I NKG2D chimera 10^5^ effector cells were co-cultured with 2 × 10^5^ target cells overnight, in presence or absence of 100 μM pamidronate. Introduction of the different chimeras in Jurkat-76 cells did not increase the percentage of CD69+ cells compared to cells transduced with γδTCR-Cl5 alone when they were co-culture together with HL60 (results comparable to no target condition) and Daudi (Fig. 4). However, an increase in CD69 positive cells was observed when cells co-transduced with γδTCR-Cl5 and NKG2D-CD28_wt_ were co-cultured together with K562, suggesting that this additional effect provided by the chimera only occurs in presence of NKG2D ligands in combination with TCR activation, as no increase was observed against other cell lines like Daudi, which expresses low NKG2D ligand levels or HL60 which despite high levels of NKG2D ligands is not recognized by γδTCR-Cl5. However, we could not exclude at this stage that increased activity is a consequence of increased expression or an altered signaling between different constructs

**Figure 4.**
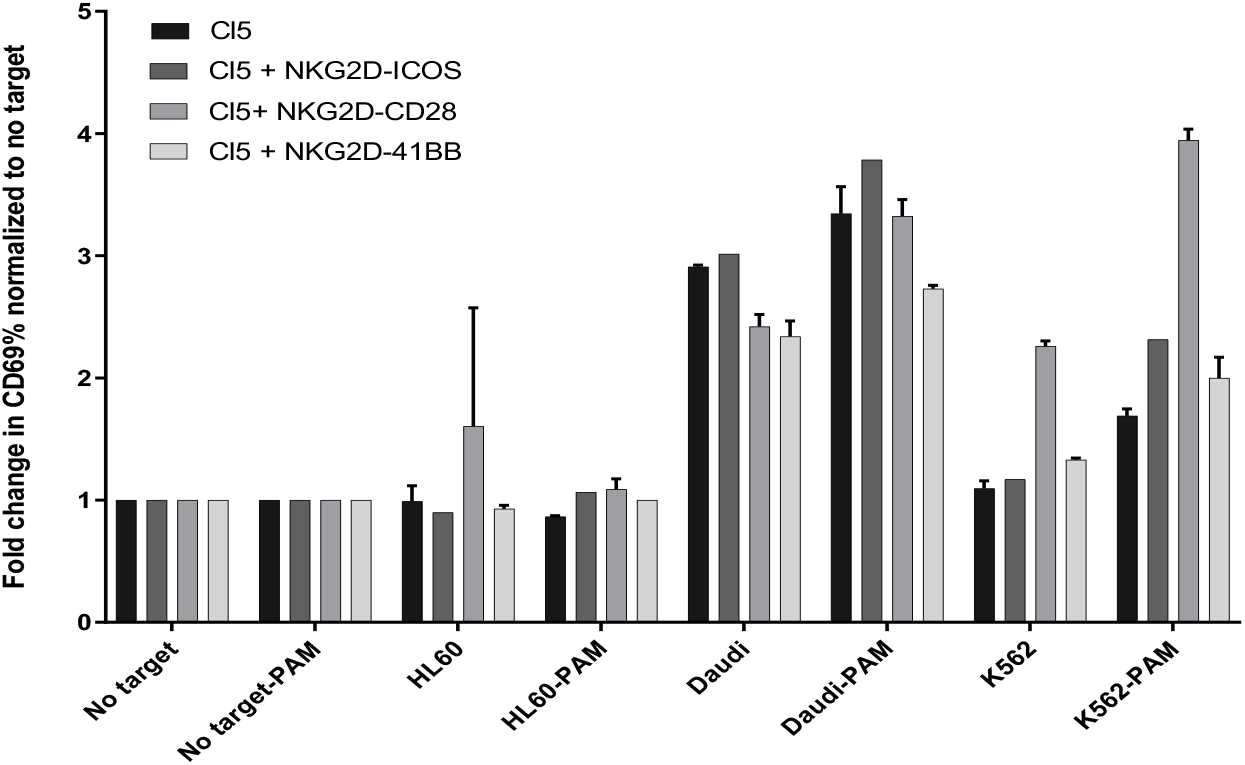
CD69 expression in transduced Jurkat-76. Percentage of CD69 positive cells was assessed by flow cytometry in Jurkat-76 cells transduced with gdTCR-Cl5 alone or gdTCR-Cl5 and the different type I chimeras after co-culturing with K562, RPMI 8226 and HL60.

### Equal expression of the chimeras by introducing CD28 transmembrane and linker domains

To assess whether differences in expression of type I NKG2D chimera is a general phenomena and to compare expression profile to engineered NKG2D_wt_, primary PBMC were engineered with γ9δ2TCR, NKG2D_wt_ and type I variants. In order to avoid that unequal expression is a consequence of different transduction efficacy of separate constructs, NKG2D_wt_ or three type I NKG2D chimeras were cloned together with the γδTCR-Cl5 into the clinical vector pmp71 (Figure 5A). PBMCs were transduced using these constructs. Expression of the three chimeric receptors and γδTCR-Cl5 was assessed by FACs (Fig. 5B). Introduction of exogenous NKG2D_wt_ further increased natural NKG2D expression as well as introduction of type I NKG2D chimera. However, again the expression level of the NKG2D-CD28_wt_ chimera was stronger when compared to the other two chimeras and NKG2D_wt_.

**Figure 5.**
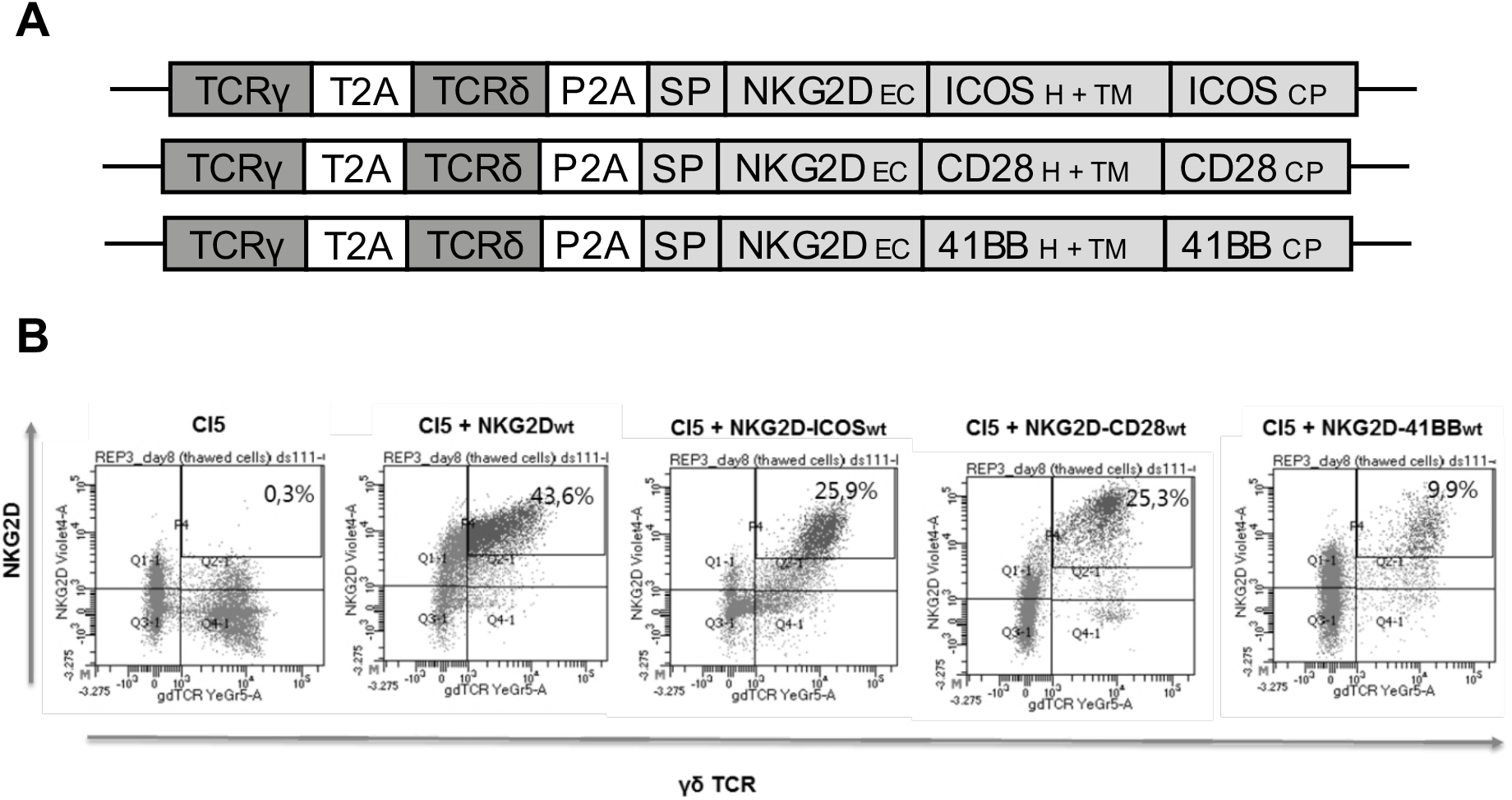
Unequal expression of type I chimeras in PBMCs. (**A**) Schematic overview of three different type I chimeras and gdTCR transgene cassette in the retroviral vector pMP71. TCRδ chain was derived from clone 5 (d) and TCRy from clone G115 (g) and F2A (derived from the foot-and- mouth disease virus) and T2A (derived from the Thosea asigna virus) refer to two different 2A ribosomal skipping sequences. (**B**) Surface expression of gdTCR-Cl5 and type I NKG2D chimeric co-receptors in Jurkat-76 cells assessed by flow cytometry.

Strength of membrane expression of proteins can depend on their transmembrane domain [13]. NKG2D-CD28_wt_ chimera expression was a consequence of its superior transmembrane domain. Therefore we next swapped the transmembrane and linker domains of the NKG2D-ICOS_wt_ and NKG2D-4-1BB_wt_ for the transmembrane and linker domains of the NKG2D-CD28_wt_ chimera (Figure 6A). The new chimeras were cloned together with the γδTCR-Cl5 or γδTCR-LM1(mock) into the clinical vector pmp71 (Figure 6B). Again PBMCs were transduced using these constructs. γδTCR-LM1 and γδTCR-Cl5 with NKG2D_wt_ or NKG2D-CD28_wt_ chimera were also taken along as controls. Expression of the three chimeras and γδTCR-Cl5 or γδTCR-LM1 was assessed by FACs (Figure 6C). The expression of the new chimeras (NKG2D-ICOS_CD28TM_ and NKG2D-4-1BB_CD28TM_) was now improved by replacing the transmembrane and linker domains by those of CD28, and it was equal to the NKG2D-CD28_wt_ chimera. In all cases, the expression of the chimeras was better compared to NKG2D_wt_.

**Figure 6.**
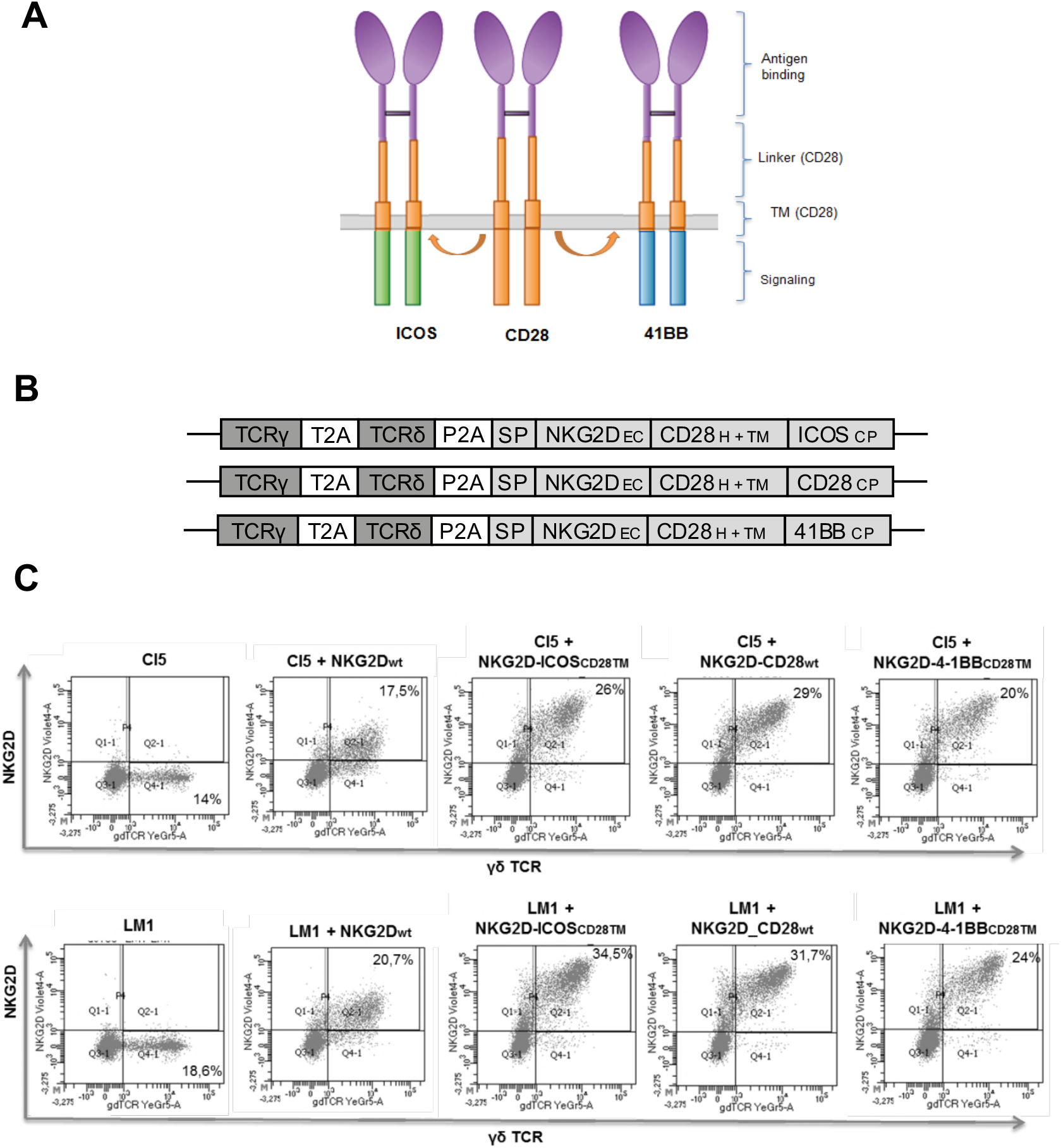
Introduction of CD28 transmembrane and linker domains increases the expression of type I designs of NKG2D-ICOS and NKG2D-4-1BB chimeric co-receptors. (**A**) Schematic diagram of the new type I chimeras containing CD28 transmembrane and linker domains (**B**) Schematic overview of new type I chimeras generated and gdTCR transgene cassette in the retroviral vector pMP71. (**C**) Surface expression of gdTCR-Cl5 and new NKG2D chimeric co-receptors in Jurkat-76 cells assessed by flow cytometry.

### Introduction of NKG2D-41BB increases cytokine release by TEG001 in presence of NKG2D ligands

In order to assess whether adding exogenous NKG2D_wt_ to TEGs increases activity of TEGs or whether adding type I chimeras would be superior, we performed an IFNγ release assay. As the NKG2D receptor is naturally expressed by many CD8+ αβT cells, CD4+ αβT cells were used as effector cells. CD4+ transduced cells were co-cultured with K562 (high levels of NKG2D ligands and recognized by γδTCR-Cl5), Daudi (low levels of NKG2D ligands but recognized by γδTCR-Cl5) and HL60 (high levels of NKG2D ligands but not recognized by γδTCR-Cl5). Pamidronate was used at different concentrations to increase the levels of phosphoantigens in the target cells and therefore increase the recognition of these cells by γδTCR-Cl5.

The most prominent change was observed for cells co-transduced with γδTCR-Cl5 and the NKG2D-4-1BB_CD28TM_ which secreted higher levels of IFNg against NKG2D-ligand high expression tumor cells K562 when compared with the TEGs that only expressed γδTCR-Cl5 (Figure 7). In addition, in this particular combination of TEGs with NKG2D-4-1BB_CD28TM_, engineered immune cells were also able to recognize NKG2D high expressing tumor cells K562 at lower pamidronate concentrations when compared to all other conditions. This increase was not observed when T cells were co-transduced with a non-functional γδTCR (LM1). Furthermore, this increase in IFNg release was not observed against Daudi (low levels of NKG2D ligands) or HL60 (not recognized by γδTCR-Cl5), suggesting that both signals (TCR activation and NKG2D ligands) are needed to induce activation of the chimera, and which is in line with the results obtained using Jurkats. This emphasized also the need to compare engineered with equivalent receptor expression, as this signal was missed in Jurkat cells when using NKG2D domains with 4-1-BB transmembrane and signaling domain. We could also not rule out that enhanced NKG2D_wt_ expression might have similar effects, however in the wt design the expression level was limited and always inferior when compared to chimeras using CD28 transmembrane domains.

**Figure 7.**
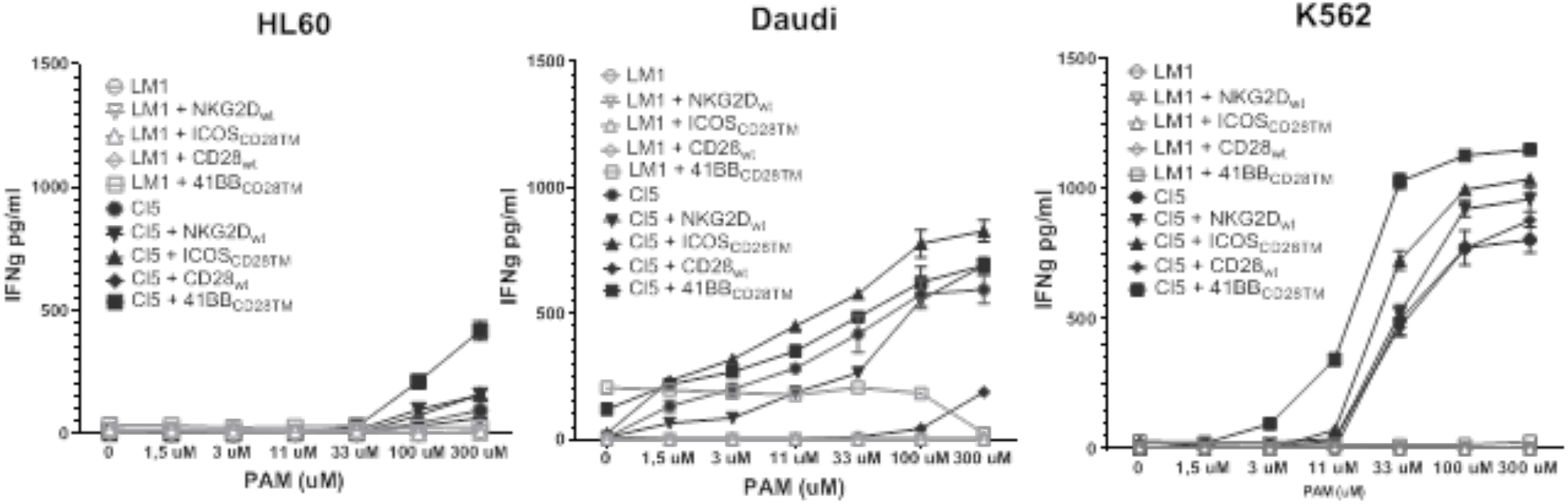
Introduction of NKG2D-4-1BB chimeric co-receptor increases IFNγ release by TEG001. Transduced T cells were incubated with K562, HL60 or Daudi at several pamidronate concentrations. After 18 hours, supernatants were harvested and analyzed for IFNγ secretion by ELISA.

### Introduction of NKG2D-41BB and NKG2D-CD28 increases killing and proliferation ability of TEG001

Next, we aimed to asses whether the NKG2D type I chimeras with the CD28 transmembrane domain were able to increase the killing ability of TEGs. Therefore, T cells transduced with γδTCR-Cl5 alone, γδTCR-Cl5 and NKG2D_wt_ or γδTCR-Cl5 and the different type I chimeras with the CD28 transmembrane domain were co-cultured with RPMI-8226 tumor cells expressing luciferase, in presence or absence of pamidronate. Co-culture was made using different effector-target ratios. After incubation, luciferin was added to the wells and signal was measured.

A mixture of CD8^+^ (75%) and CD4^+^ (25%) αβ T cells co-transduced with γδTCR-Cl5 and the chimeras NKG2D-CD28_wt_ and NKG2D-4-1BB_CD28TM_ showed increased killing ability compared to γδTCR-Cl5 alone (Fig. 8). This increase in killing ability was especially remarkable at lower E:T ratios and was not observed in T cells co-transduced with mock γδTCR (LM1) and the different NKG2D chimeras. Furthermore, cells co-transduced with γδTCR-Cl5 and NKG2D wt also showed an increment in killing compared to γδTCR-Cl5 alone. However, also an increase in specific lysis was observed when NKG2D wt was co-transduced with γδTCR-LM1 (mock).

**Figure 8.**
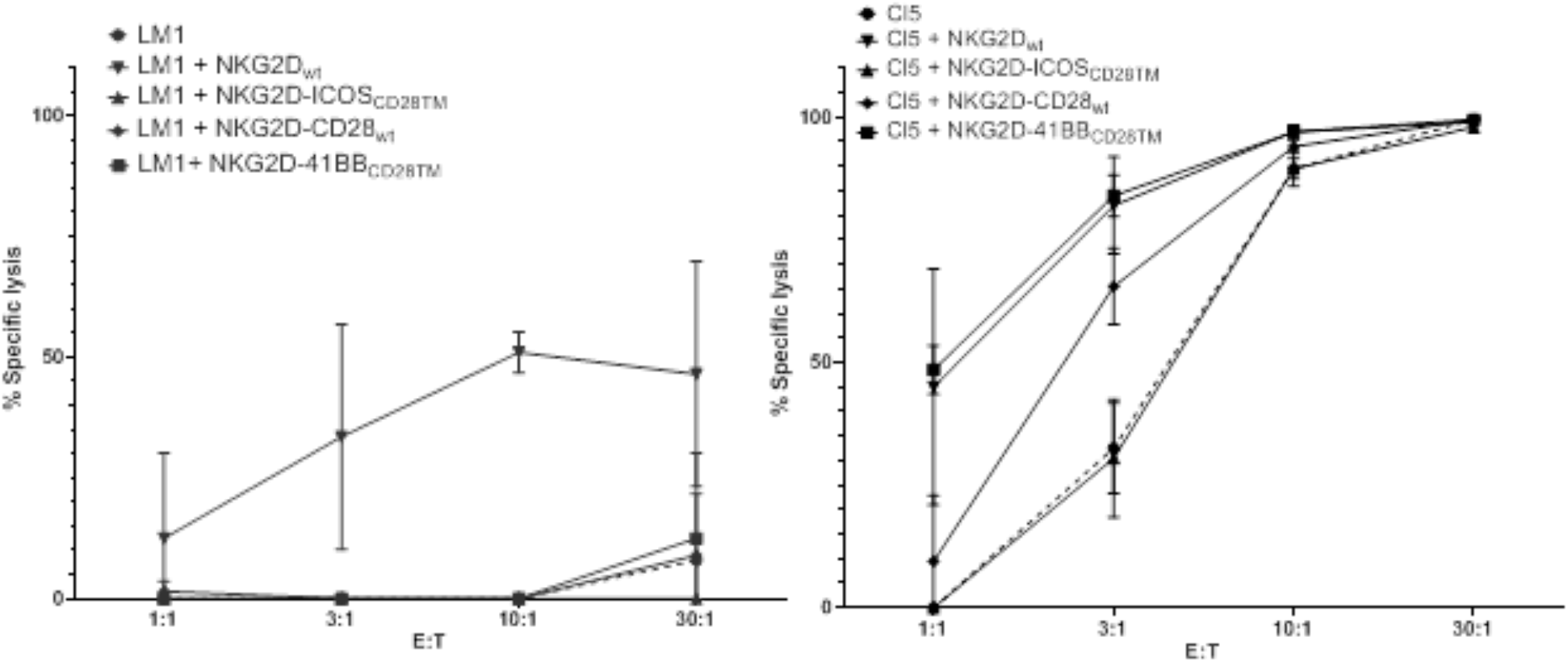
Introduction of NKG2D-4-1BB chimeric co-receptor increases killing ability of TEG001. Transduced T cells were incubated with RPMI 8226-luc+ cells in presence of 10 μM PAM. Specific lysis was calculated using the formula % specific lysis = 100 × [(experimental data − spontaneous cell death)/(maximum cell death − spontaneous cell death)]. When this calculation provided a negative value, 0% was assigned as the result.

### Introduction of NKG2D chimeras improves proliferation capacity of TEG001

Different co-stimulatory domains have been reported to mainly affect proliferation capacity. Therefore we examined proliferation capacity of TEGs transduced with NKG2D_wt_ and the different type I NKG2D chimeras with CD28 transmembrane domains by using a dye dilution approach. Again CD4+ αβT cells selected cells were used as it allowed also to assess the additional impact of NKG2D_wt_. They were fluorescently labeled using Cell Trace violet dye and co-cultured either with HL60 (NKG2D ligand negative) or RPMI-8226 (NKG2D ligand positive) tumor cells in presence of 100 μM pamidronate. After 6 days, cells were analyzed by flow cytometry. As expected, cells transduced with γδTCR-LM1 (mock) were not able to proliferate when co-cultured together with tumor cells (Figure 9). Moreover, the introduction of NKG2D_wt_ did not change the proliferation rate compared to T cells transduced with γδTCR-Cl5 alone or when NKG2D_wt_ was expressed. In contrast TEGs coexpressing three different type I chimeras with the CD28 transmembrane domain showed an improved proliferation capacity as indicated by the lower CTV intensity. Furthermore, this increase in proliferation was only observed when T cells were co-cultured together with RPMI 8226, but not with HL60. As both, RPMI 8226 and HL60 are expressing high levels of NKG2D ligands, but only RPMI-8226 is recognized by γδTCR-Cl5, these data suggest that γ9δ2TCR activation is essential for any additive effect of the chimeras.

**Figure 9.**
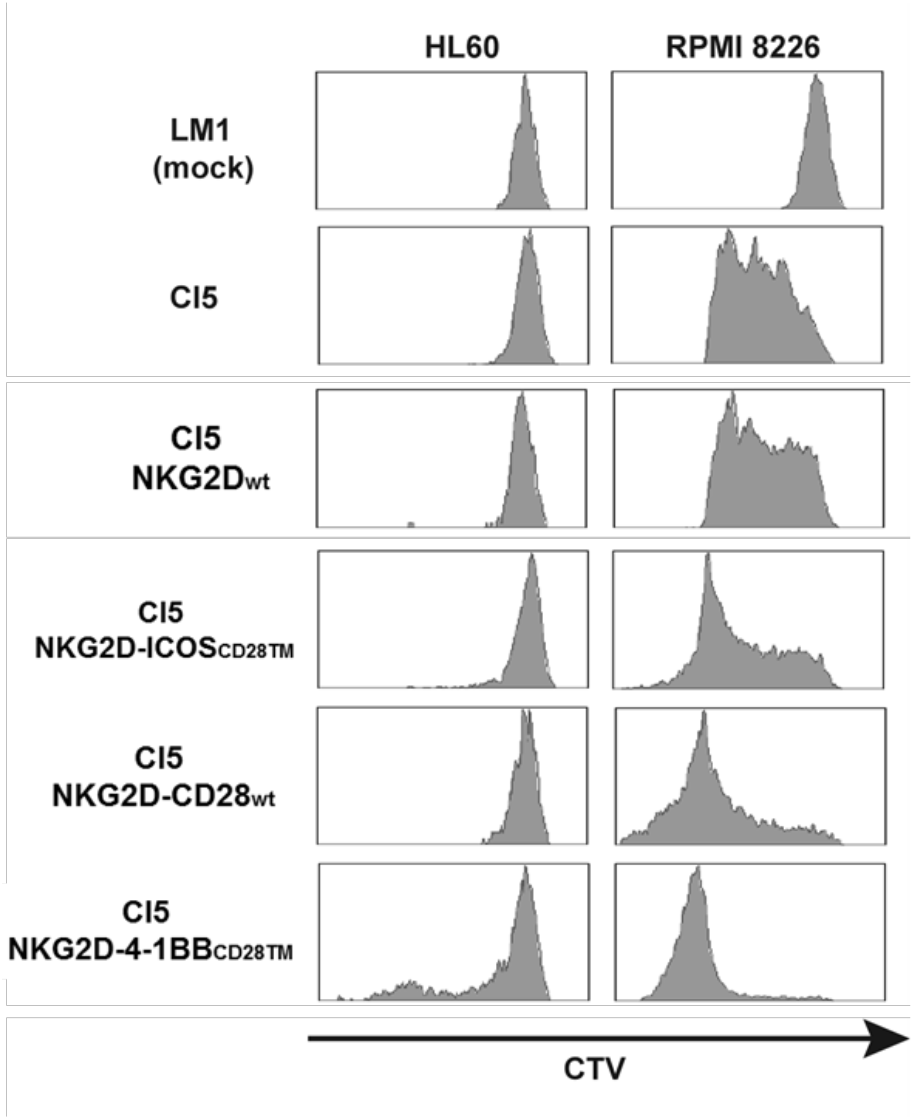
Introduction of NKG2D chimeric co-receptors improves proliferation ability of TEG001. Transduced T cells were incubated either with HL60 or RPMI 8226 cells in presence of 100 μM PAM. On day 6 CTV intensity was assessed by flow cytometry.

## Discussion

Different chimeras were designed in this project, by fusing the extracellular domain of NKG2D to the cytoplasmic domain of three different costimulatory proteins (ICOS, CD28, 4-1BB). According to the orientation of the proteins, type II and type I chimeras were designed. Only chimeras with a type I orientation were expressed. However, expression between type I chimeras was unequal when the transmembrane and linker domains of the different costimulatory proteins were used, with NKG2D-CD28_wt_ chimera always be the best expressed when compared to NKG2D-ICOS_wt_ and NKG2D-4-1BB_wt_. To improve expression of other chimeras, swapping the transmembrane and linker domains of NKG2D-ICOS and NKG2D-4-1BB by NKG2D-CD28’s domains was explored (NKG2D-ICOS_CD28TM_ and NKG2D-4-1BB_CD28TM_). This modification increased the expression of both chimeras and equaled the expression between all of the chimeras.

Cells co-transduced with γδTCR-Cl5 and the three chimeras were tested in different functional assays and compared with cells transduced with γδTCR-Cl5 alone (TEG001). Altogether, these results suggest that the introduction of NKG2D-CD28_wt_ and NKG2D-4-1BB_CD28TM_ increase the activity of TEG001.

---

**Supplementary figure 1.**
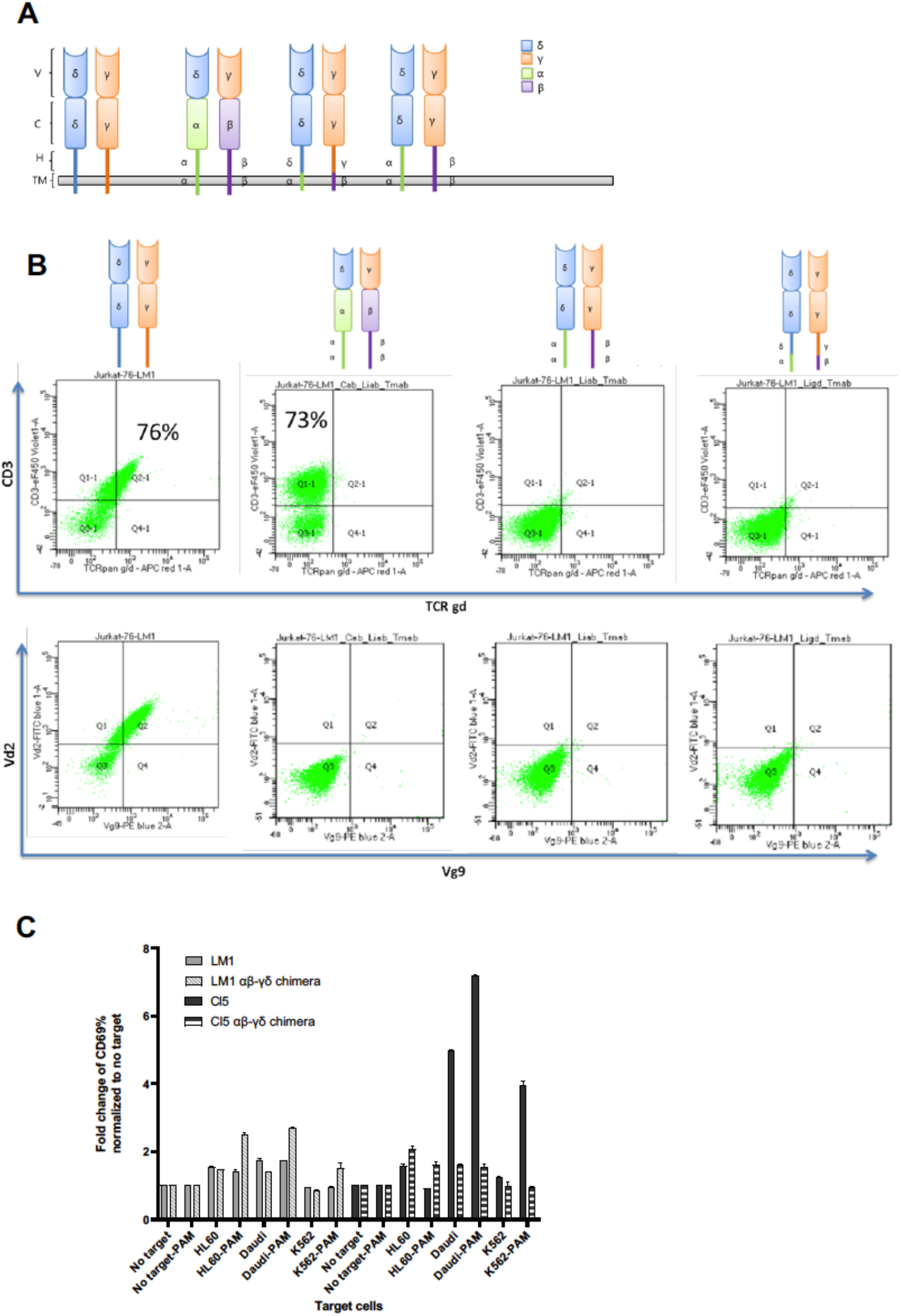
First version of a αβ-γδ TCR chimeras. **(A)** Schematic overview of three different αβ-γδ TCR chimeras. (V) variable, (C) constant, (H) hinge and (TM) transmembrane domains from αβ or γδ TCRs were used **(B)** Surface expression of αβ-γδ TCR chimeras in Jurkat-76 cells assessed by flow cytometry. **(C)** Percentage of CD69 positive cells was assessed by flow cytometry in Jurkat-76 cells transduced with gdTCR-Cl5, gdTCR-LM1 (mock) or dfferent αβ-γδ TCR chimeras after co-culturing alone or with K562, Daudi and HL60.

**Supplementary figure 2.**
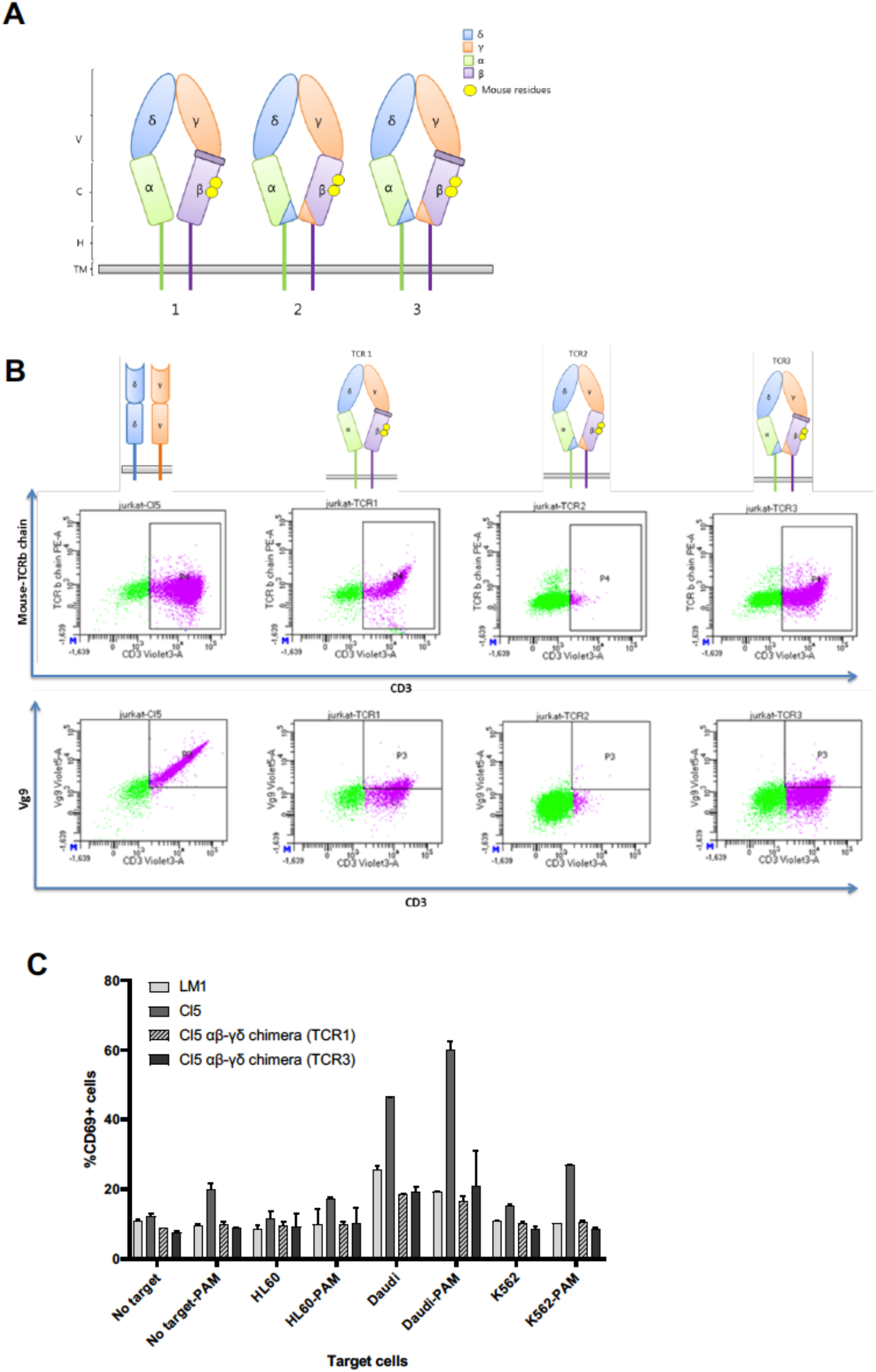
Second version of αβ-γδ TCR chimeras. **(A)** Schematic overview of the second version of three different αβ-γδ TCR chimeras. (V) variable, (C) constant, (H) hinge and (TM) transmembrane domains from αβ or γδ TCRs were used **(B)** Surface expression of the second version of αβ-γδ TCR chimeras in Jurkat-76 cells assessed by flow cytometry. **(C)** Percentage of CD69 positive cells was assessed by flow cytometry in Jurkat-76 cells transduced with gdTCR-Cl5, gdTCR-LM1 (mock) or αβ-γδ TCR chimera named as 1 and 3 after co-culturing alone or with K562, Daudi and HL60.

## Appendix

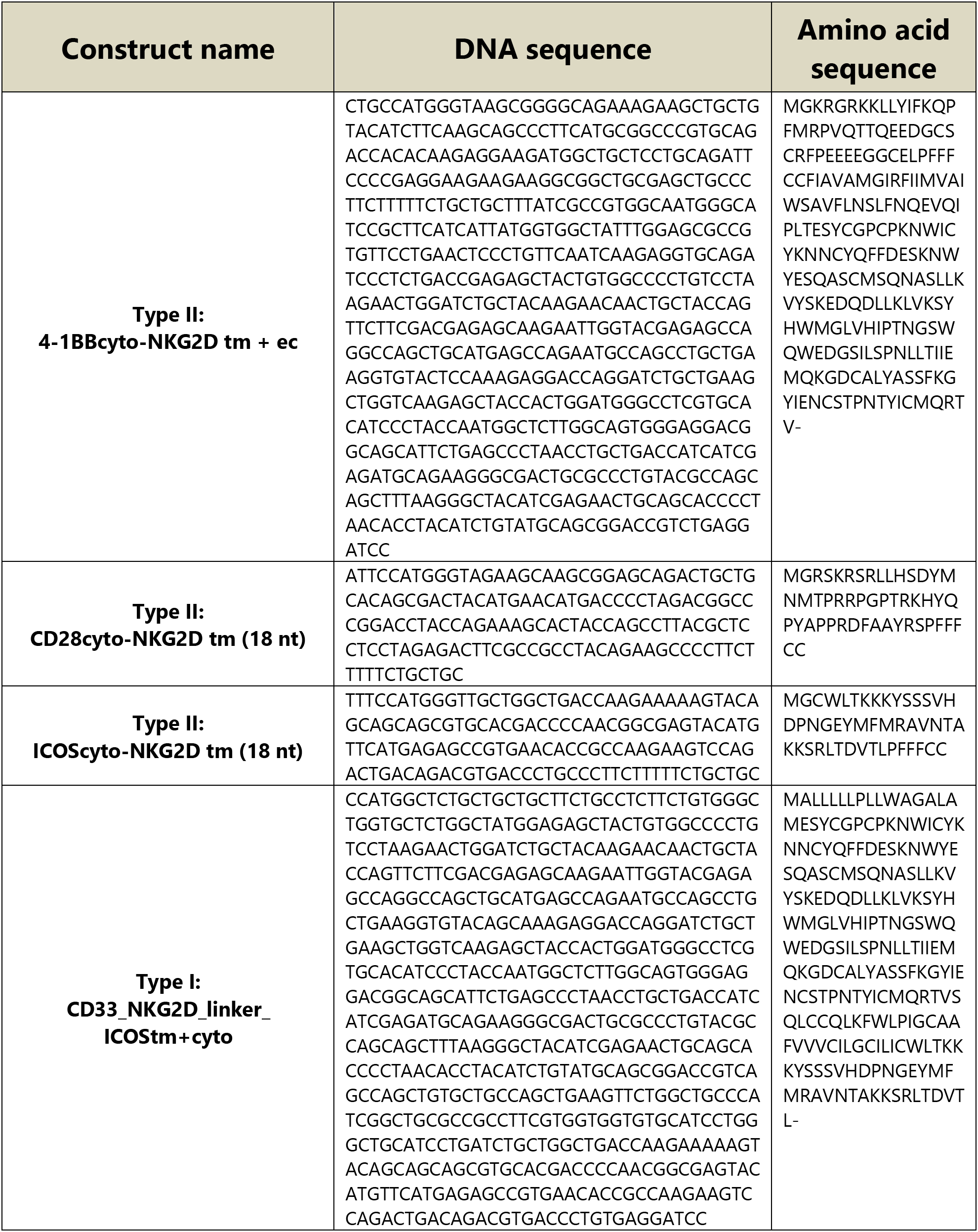
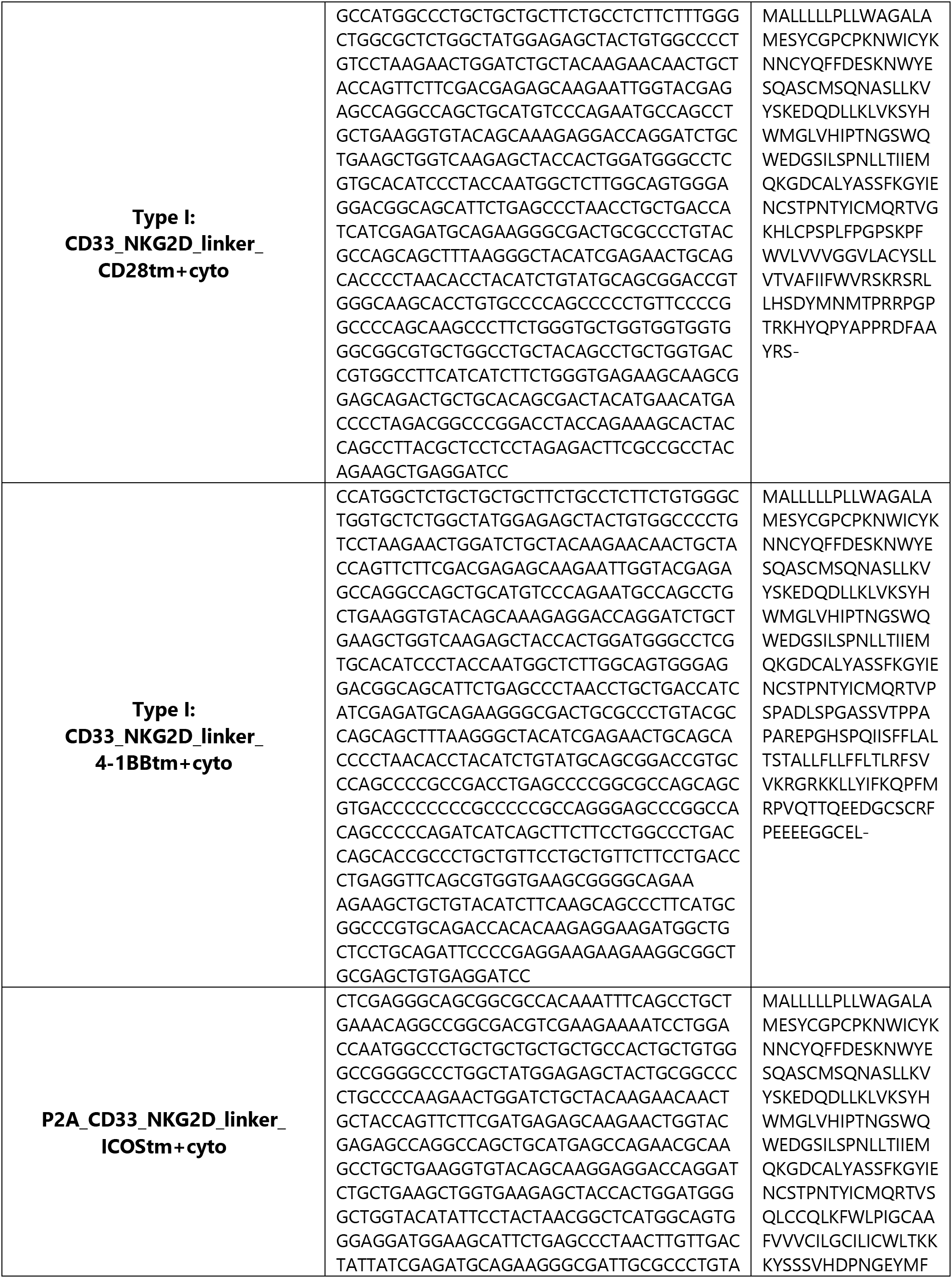
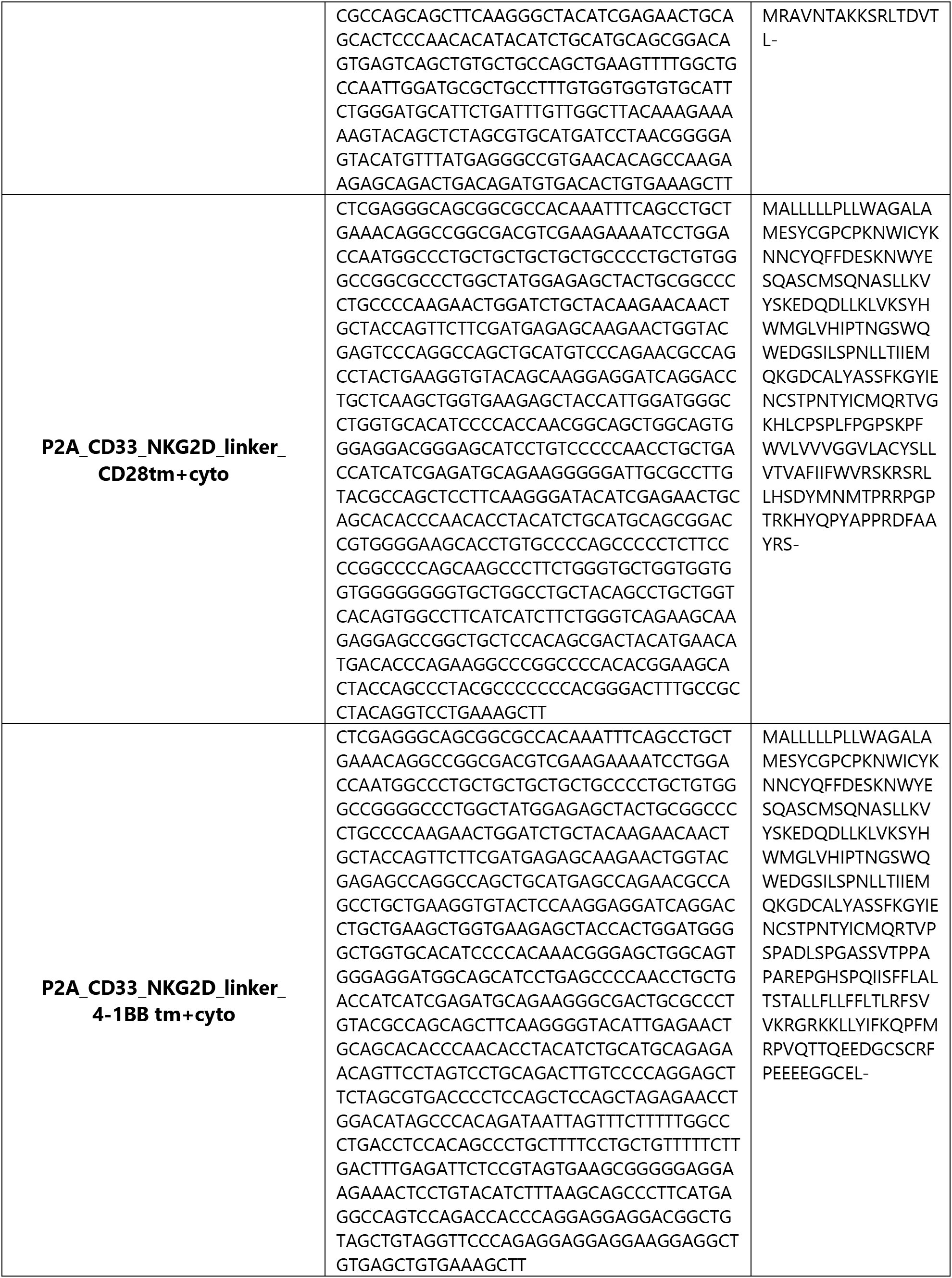
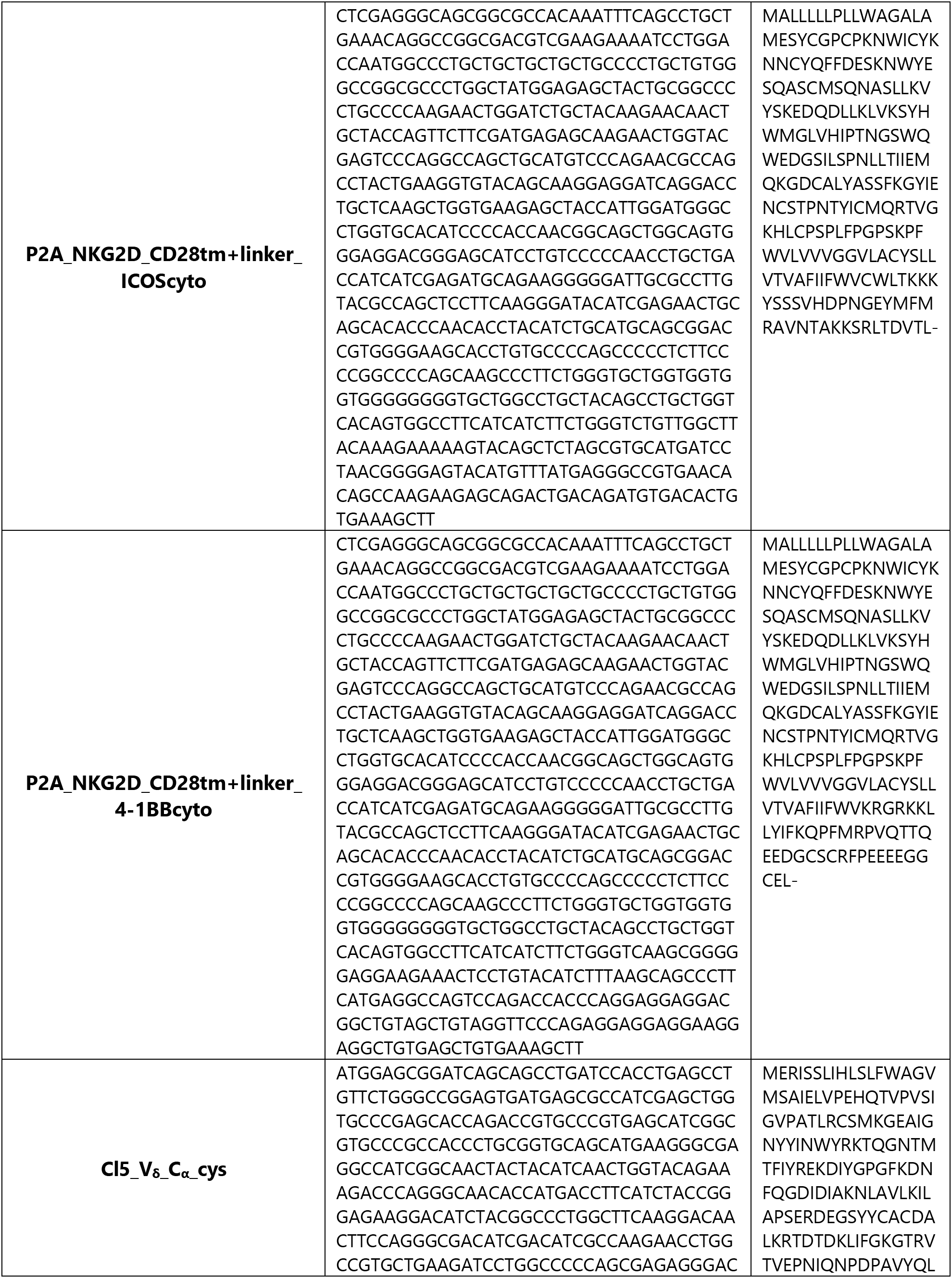
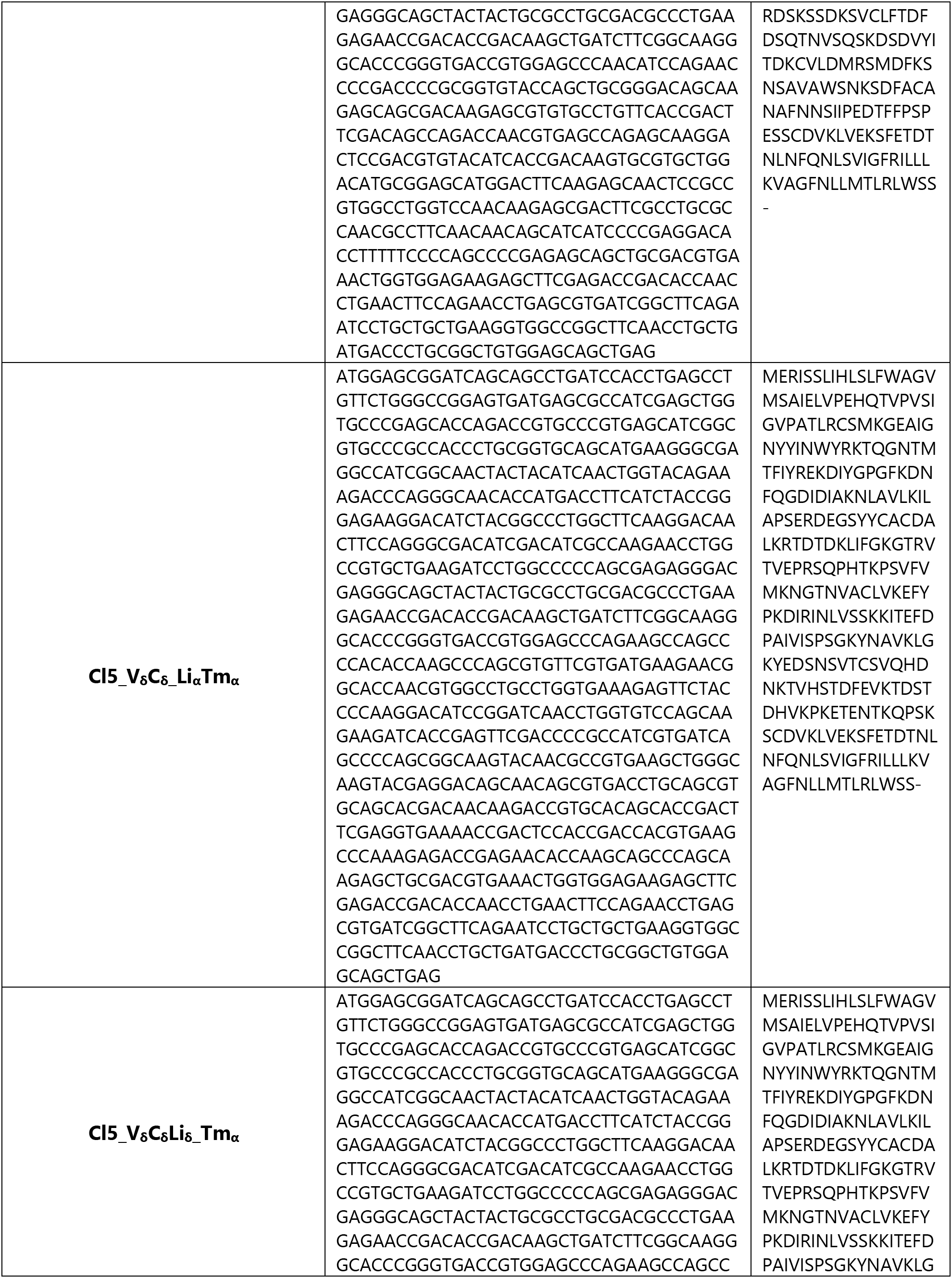
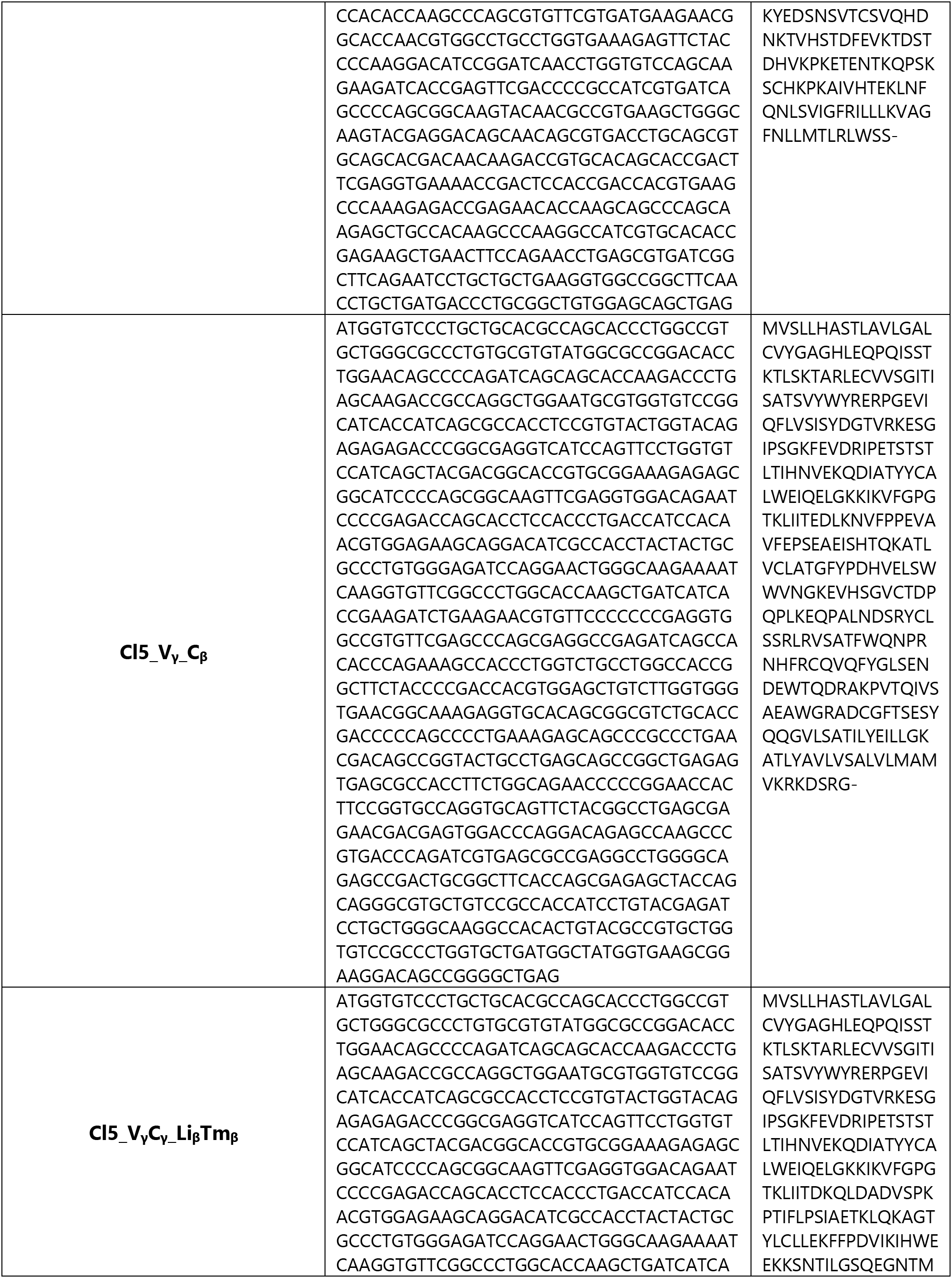
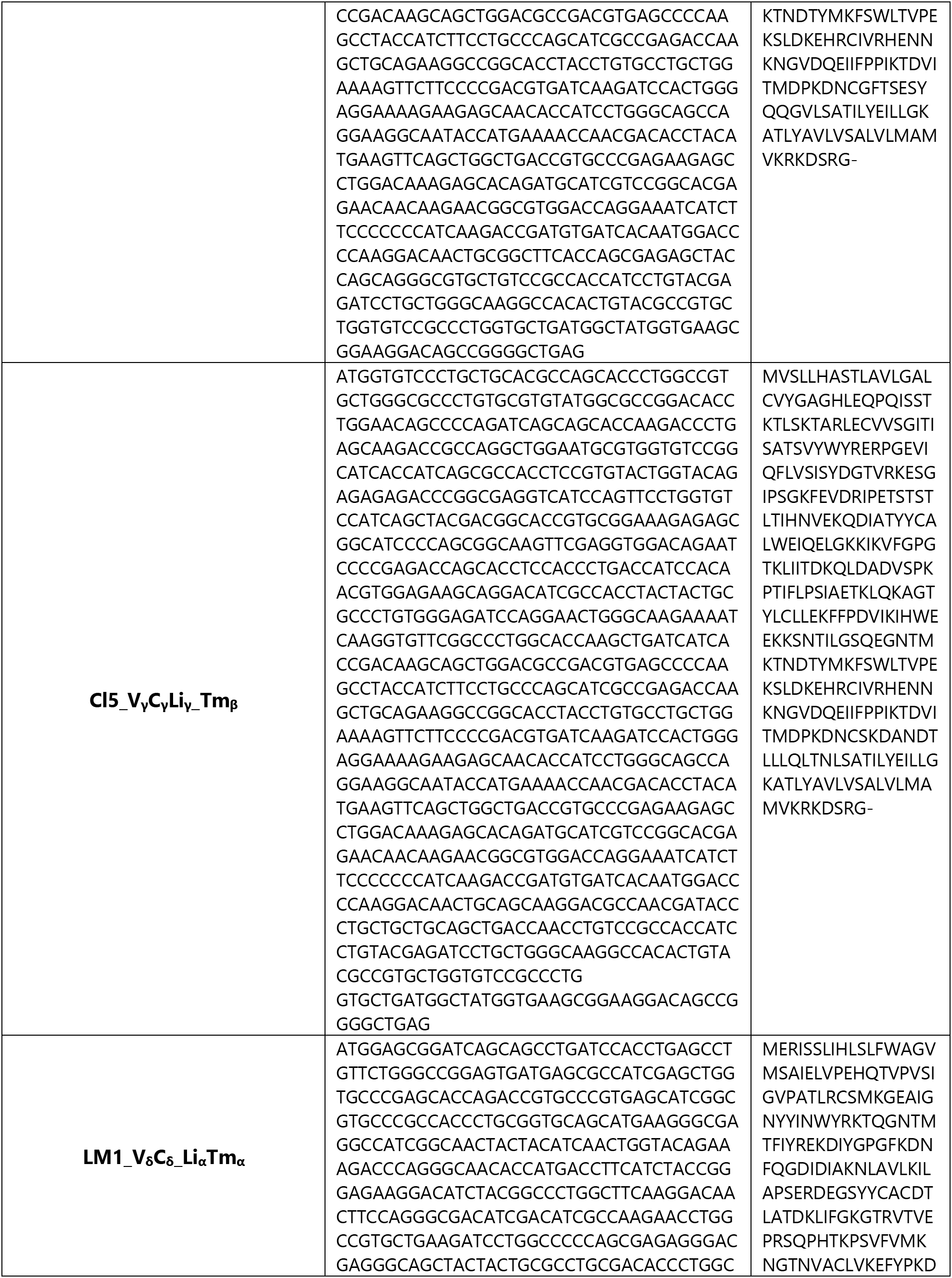
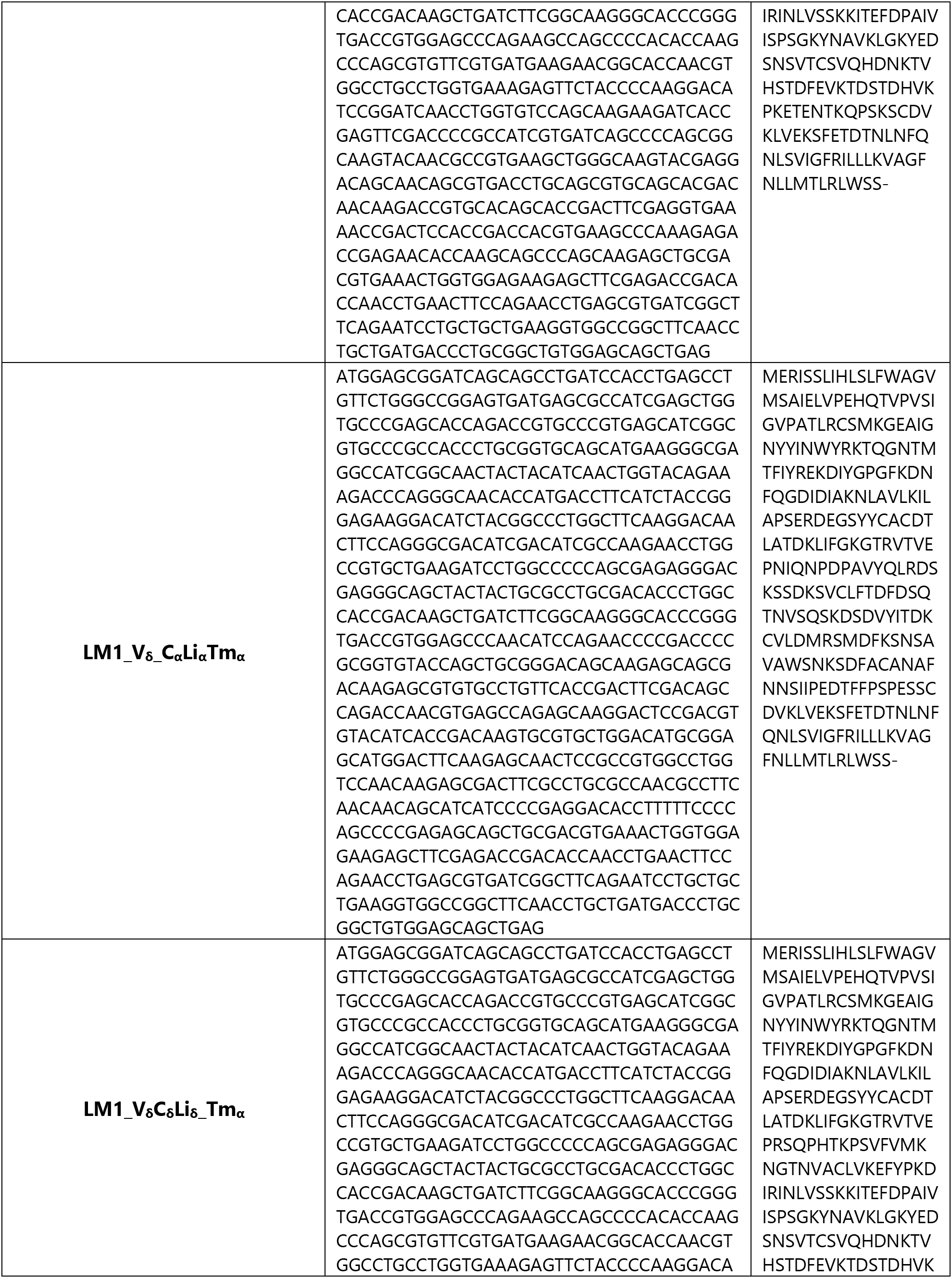
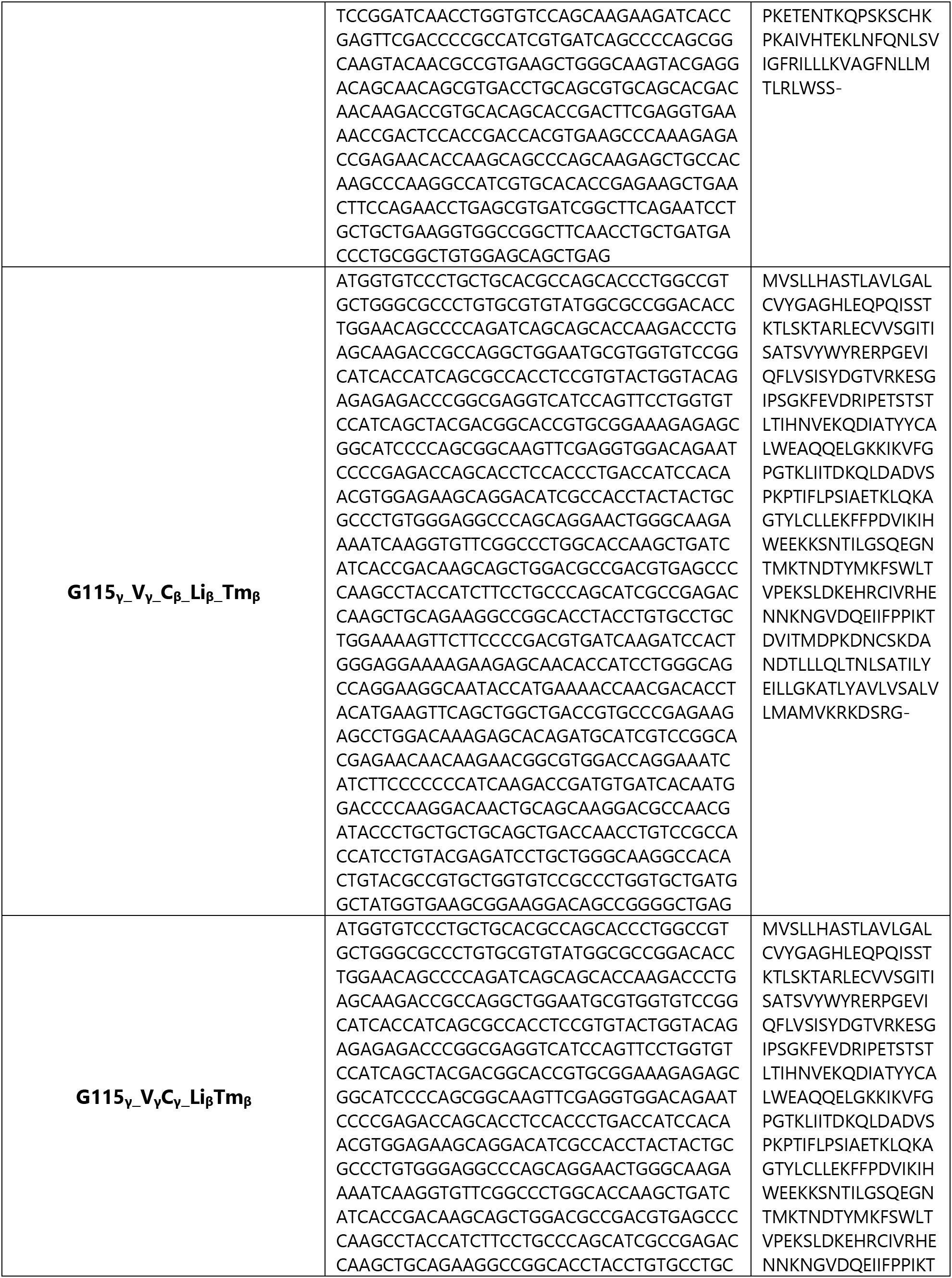
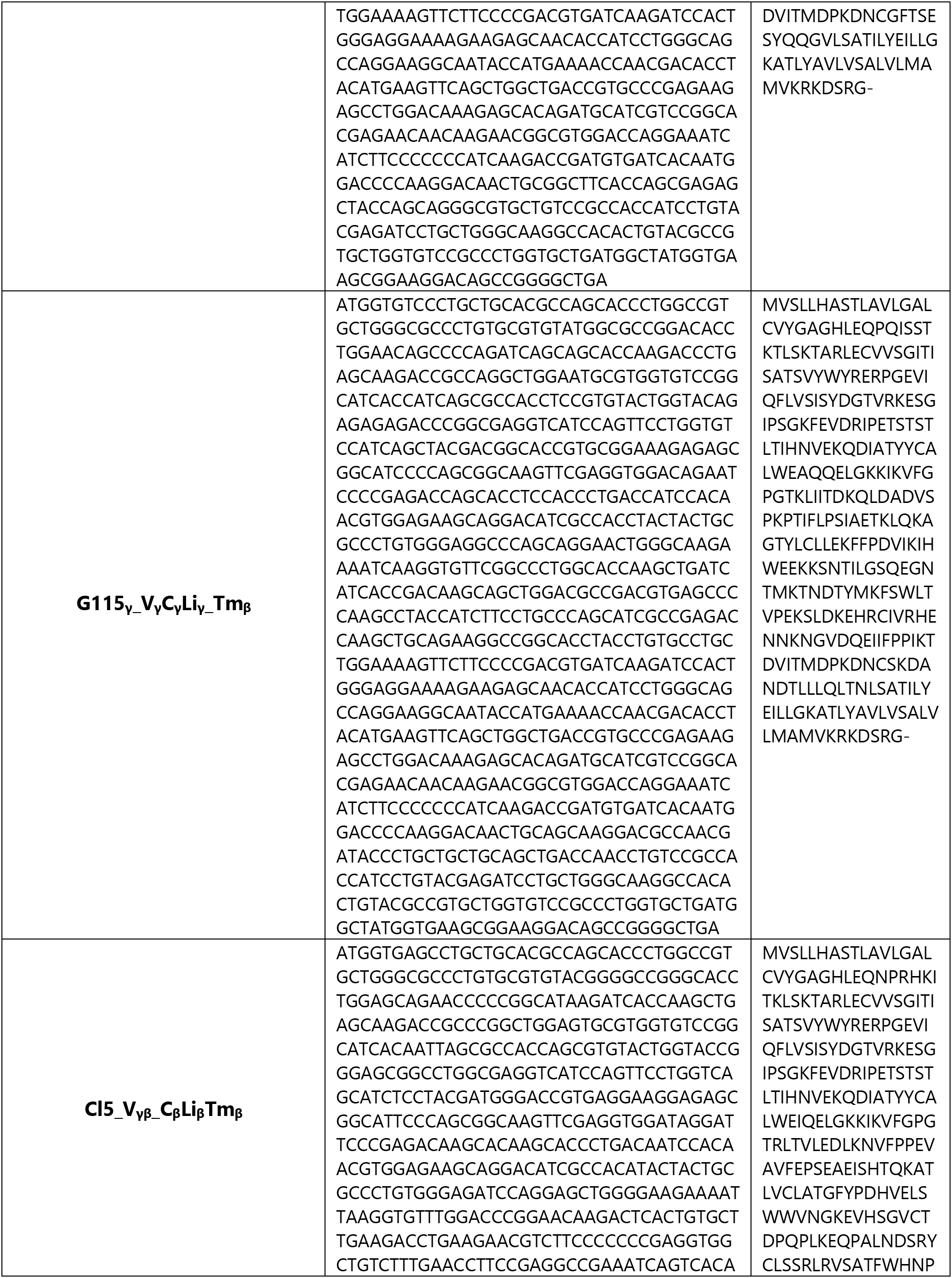
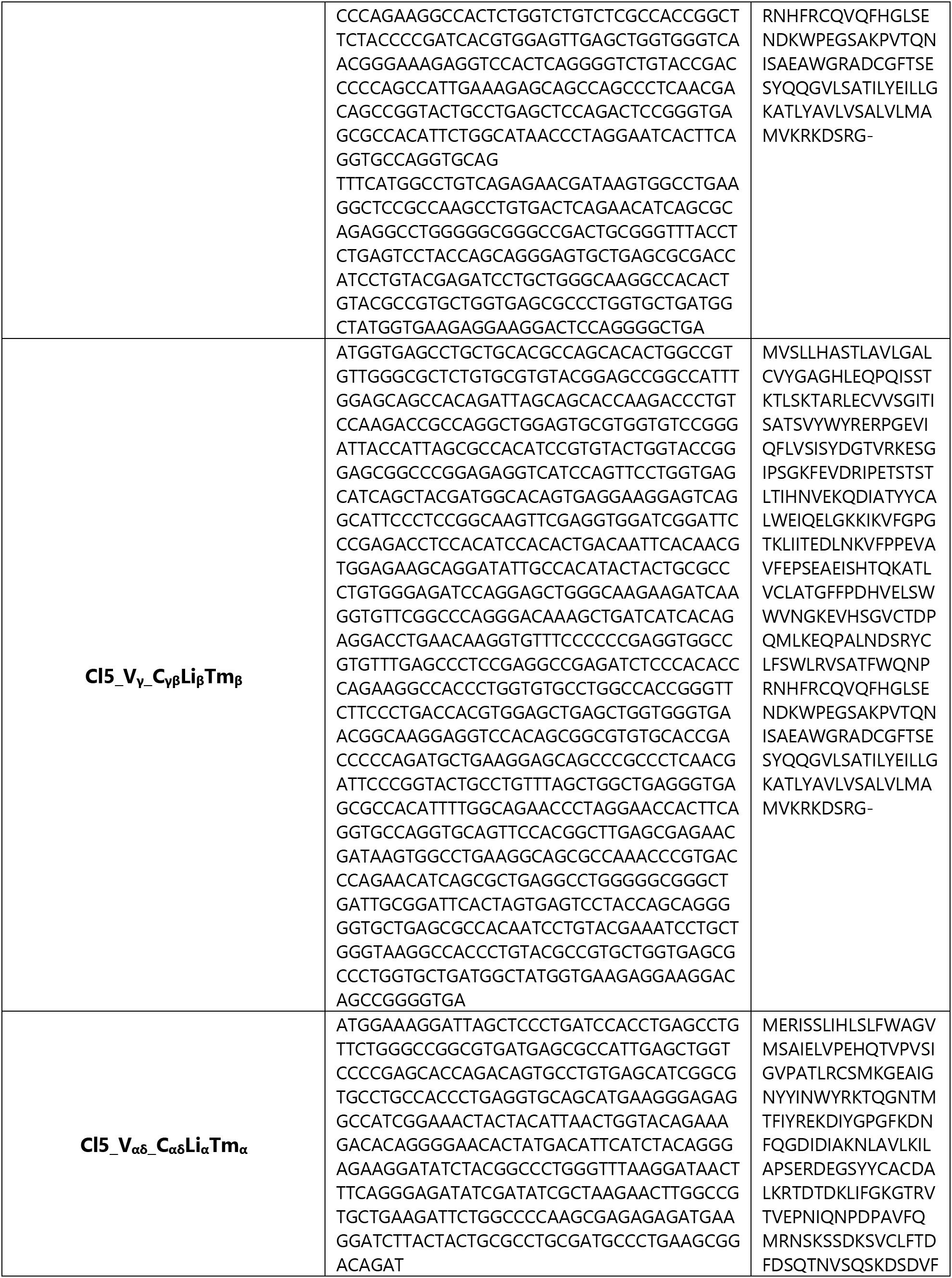
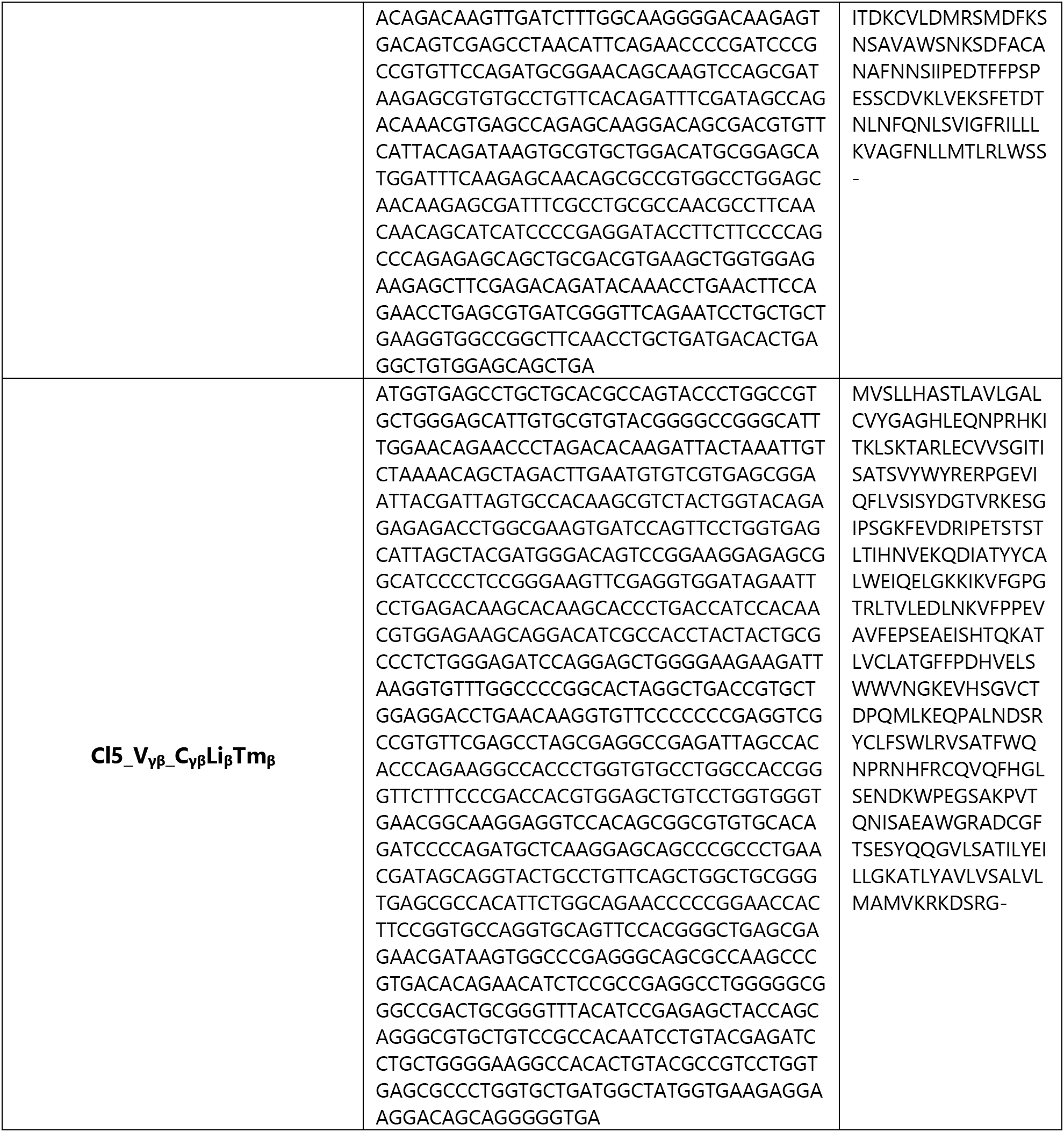

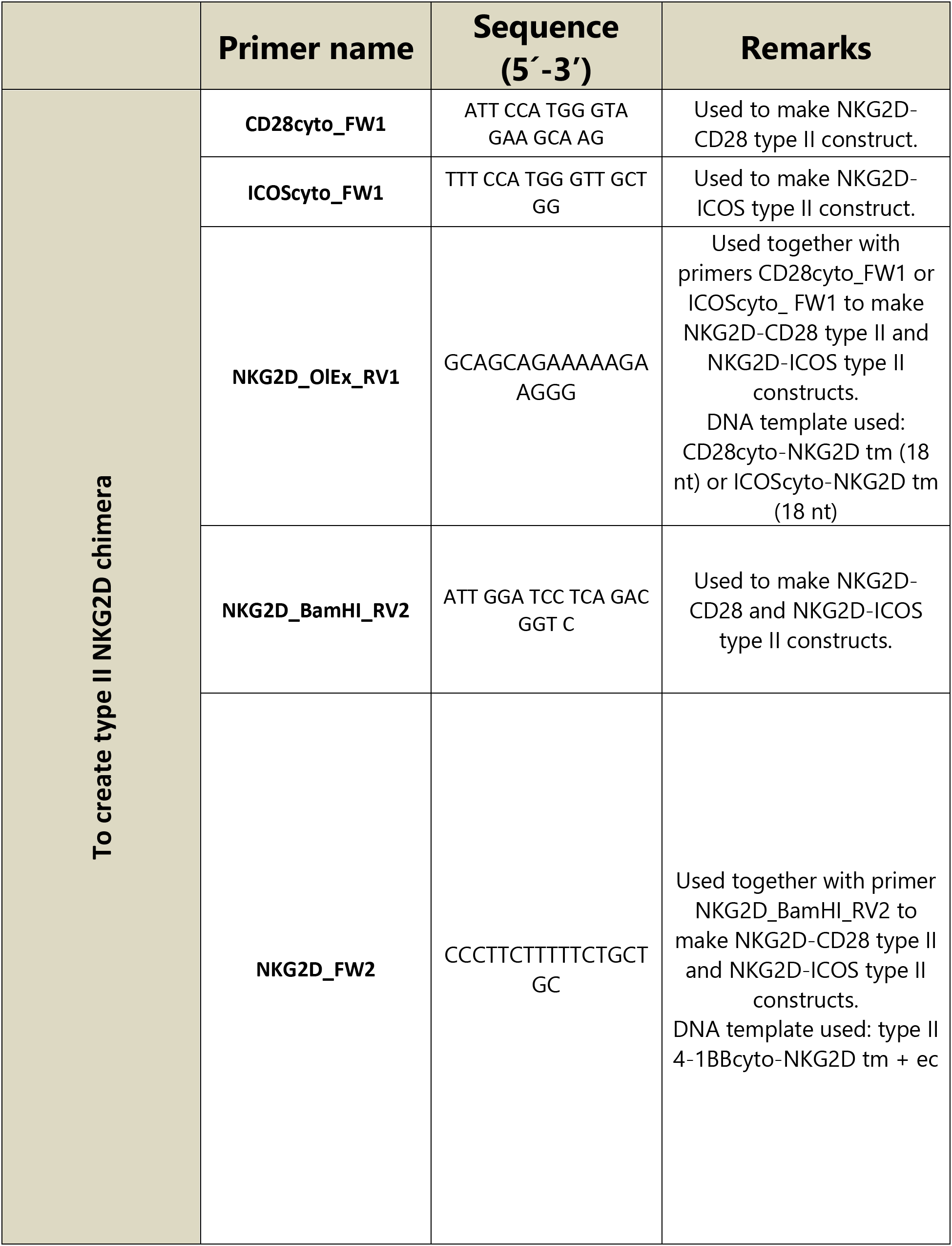
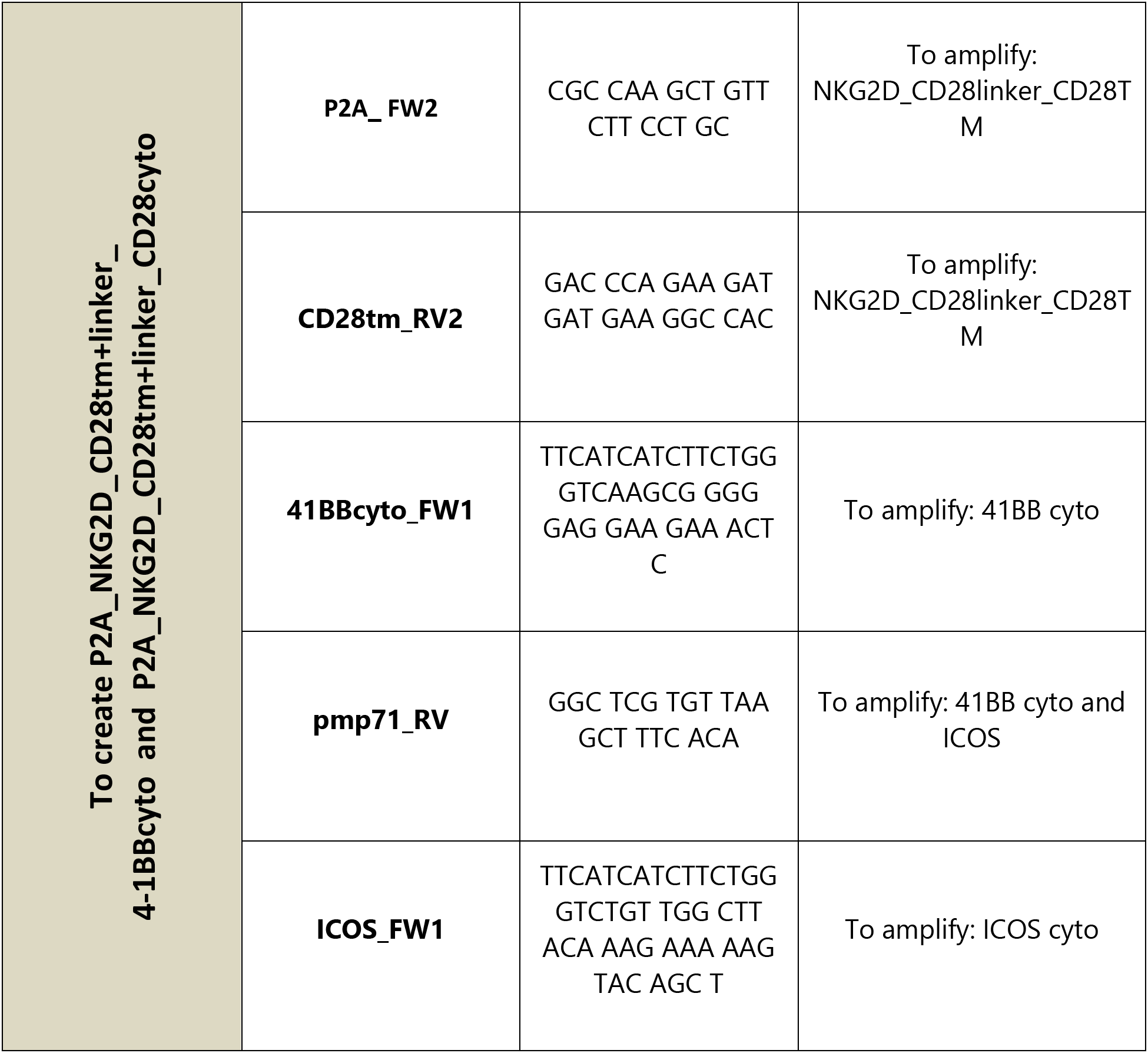

